# Genetically encoded systems for cytosolic activity recording and molecular archiving across time

**DOI:** 10.64898/2026.06.15.732182

**Authors:** Yixiao Yan, Yixuan Wang, Lirong Zheng, Jormay Lim, Nan Xiang, Changyang Linghu

## Abstract

Cellular functions are governed by dynamic changes in molecular abundance, state, activity, and interactions within the cytosol. Current molecular tape recorders leverage nuclear transcriptional programs in eukaryotic cells to provide scalable readout of gene regulation dynamics, but cytosolic activity, molecular abundance, and state remain inaccessible. Here, we introduce CyTRACE, a genetically encoded, scalable framework for cytosolic activity recording and molecular archiving across time. CyTRACE ventures beyond the cell nucleus, deploys cytosol-localized protein tape monomers as direct activity sensors and molecule catchers, and offers physiological sensitivity, spatiotemporal resolution, and programmability. Within this modular framework, CytoSense records protease and kinase signaling activities while CytoCatch archives cytosolic proteins, each with temporal continuity in single mammalian cells. CyTRACE thus opens cytosolic biology to scalable activity readout and molecular interrogation.

## 1 Introduction

Many fundamental cellular processes in health and disease are mediated by molecules in the cytosol. The abundance, degradation, modification, activity, and interaction of cytosolic molecules often exhibit complex temporal dynamics that collectively govern how cells sense, compute, and respond to their internal states and external environments across time^1–3^. Resolving these temporal dynamics at scale remains a major challenge in biomedical research and therapeutic development.

Fluorescent reporters enable real-time, spatiotemporally resolved measurements of cytosolic activities in living cells, while the spatiotemporal scale of readout is often constrained by an inherent tradeoff between spatial resolution and scale under live-imaging systems and photobleaching of fluorescent reporters. Endpoint assays provide large-scale, high-resolution readout of molecular contents and their states, but capture only one static, endpoint snapshot at a time rather than a continuous record of temporal trajectories. Activity-dependent tagging systems enable labeling of active cell populations within a defined time window prior to the endpoint^4–9^, yet are typically limited to a single tagging window per cell. Split-HaloTag-based chemical labeling approaches^10,11^ achieve direct recording of cytosolic signal activity in up to three temporally discretized windows distinguished by spectrally distinct fluorescent HaloTag ligands. However, direct quantitative comparison of recorded intensities across time windows may be challenging in practice, because the labeling intensity readout from distinct fluorescent dyes can be influenced by differences in their intrinsic brightness, spectral-channel-specific imaging conditions, and by variability in the loading of individual dyes across cells.

Nucleic acid- and protein-assembly-based molecular recorders offer alternative strategies by encoding cellular activity into durable records on molecular substrates, such as nucleic acid sequences or spatial sequences of protein monomers, for post hoc scalable readout^12–18^. These systems often leverage transcriptional regulatory elements in the cell nucleus to sense gene regulation dynamics in eukaryotic cells and control the expression of reporter modules, which subsequently write information onto the recording substrate. While transcriptional regulatory elements are powerful for capturing gene regulation dynamics, inferring cytosolic processes from their activity introduces confounds and delays from transcriptional, and sometimes translational, intermediates^19–21^. Furthermore, many cytosolic activities lack well-defined transcriptional regulatory elements capable of faithfully reporting their dynamics^22^. For example, many key signaling pathways are driven by and functionally executed through post-translational mechanisms, such as protein modifications, translocation, and turnover, rather than direct transcriptional induction^23–26^. Thus, developing molecular systems capable of direct cytosolic activity recording, and even archiving cytosolic molecules themselves, for post hoc analysis across time would unlock powerful, complementary capabilities, expanding the toolkit for studying cytosolic processes in health and disease at scale.

Here, we present CyTRACE (Cytosolic spatioTemporally Resolved Analog reCording and capturE), a genetically encoded, modular framework for recording cytosolic activities and archiving cytosolic proteins with physiological sensitivity, spatiotemporal resolution, and scalability. Building on the molecular architecture of protein tape recorders (**Fig. 1, A** and **B**)^14,15^ and going beyond their transcriptional recording capability^12–18^, CyTRACE utilizes cytosol-localized tape monomers as direct activity sensors (the “CytoSense” system, **Fig. 1, C** and **D**) and molecule catchers (the “CytoCatch” system, **Fig. 1, E** and **F**) for post hoc scalable readout and analysis of cytosolic activities and proteins. CytoSense directly senses cytosolic protease and kinase signaling activities and encodes them along the protein tape assembly for post hoc readout of their temporal dynamics. CytoCatch physically captures proteins in the cytosol and archives them along the protein tape assembly for post hoc interrogation of their temporal abundance and molecular states. CyTRACE is temporally continuous and longitudinally intensity-resolved, supporting causal inference on the activation, inhibition, or persistence of recorded activity and on the abundance and state transitions of archived molecules relative to earlier time points and prior states within individual cells. By venturing beyond the cell nucleus, the CyTRACE framework establishes a general strategy for spatiotemporally resolved recording of cytosolic activities and archiving of cellular molecules for scalable post hoc analysis across cell populations.

**Fig. 1.**
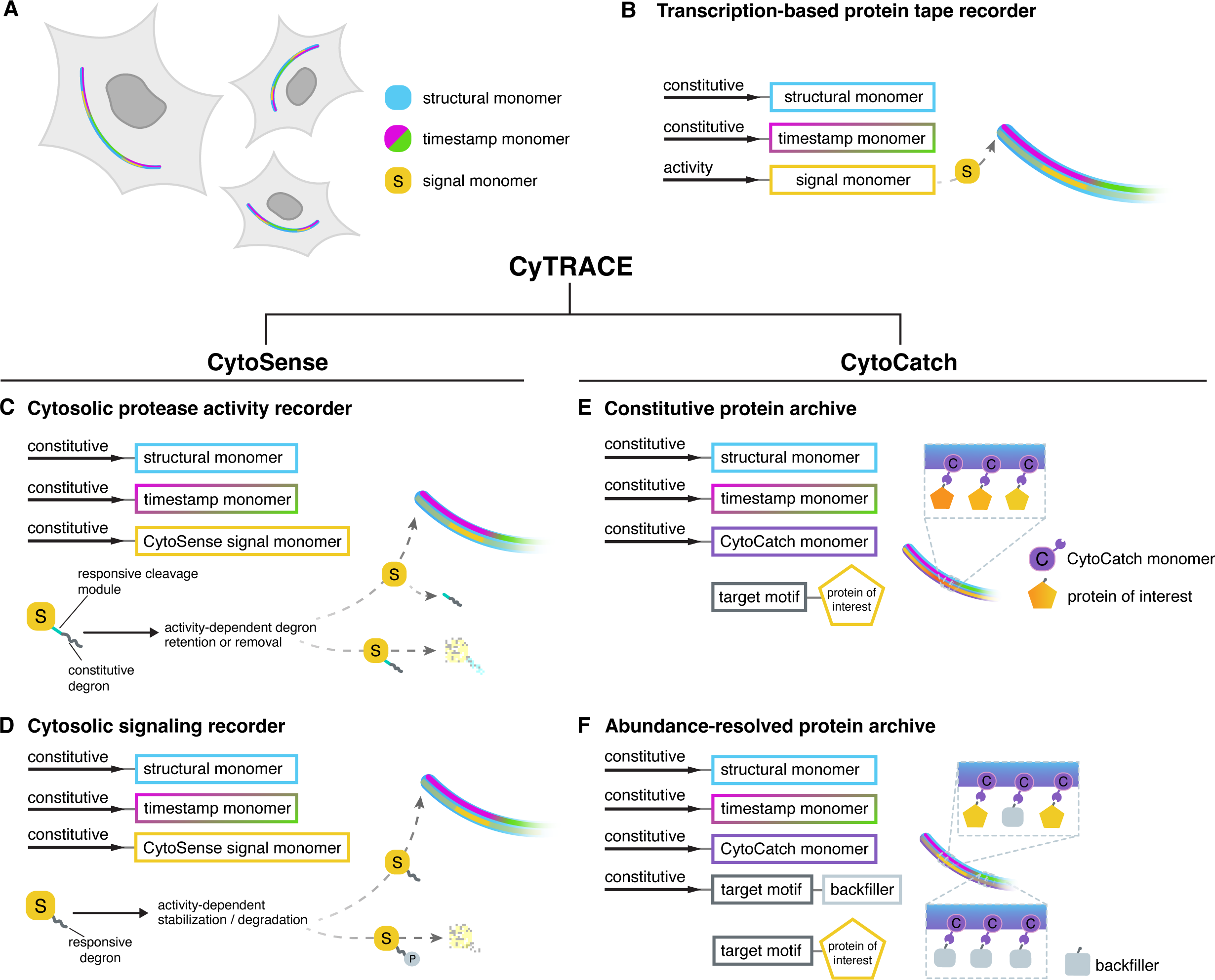
The CyTRACE framework for cytosolic activity recording and molecular capture. (**A-B**) Schematic of current protein tape recorders, using CytoTape as an example. (**A**) Intracellular protein assembly encodes temporal cellular activity along a spatial axis with three modular components that differ only by a short epitope tag: structural monomers (blue), timestamp monomers (magenta/green), and signal monomers (yellow), enabling post hoc scalable readout of temporal activity within single cells. (**B**) Existing protein tape recorders track gene regulation dynamics via transcriptionally regulated signal monomer expression, followed by subsequent signal monomer incorporation over time along an elongating protein assembly established by constitutively expressed structural monomers. (**C-F**) Building on protein tape recorders and going beyond their transcriptional recording capability, the CyTRACE framework comprises two highly programmable systems, CytoSense (**C-D**) and CytoCatch (**E-F**). The CytoSense system (**C-D**) continuously records cytosolic activity, utilizing the signal monomer of the protein tape recorder as a direct activity sensor. Cytosolic activities modulate the balance of stabilization and degradation of constitutively expressed CytoSense signal monomers, regulating their activity-dependent incorporation into the elongating protein assembly. Once incorporated, signal monomers are protected from degradation, providing a durable temporal record of signaling activity. (**C**) CytoSense records protease activity via a signal-responsive cleavage module positioned between the protein tape recorder monomer and a constitutive degron as a protease activity sensor. (**D**) CytoSense records signaling activity via a signaling-responsive degron as a signaling activity sensor. The CytoCatch system (**E-F**) continuously captures and temporally archives cytosolic proteins of interest (POIs) along the elongating protein assembly, allowing post hoc interrogation of protein abundance and molecular states across time. (**E**) CytoCatch archives POIs with a target motif along the growing protein assembly via a constitutively expressed CytoCatch monomer as a molecular catcher, which comprises a target motif-binding domain fused to the protein tape monomer. (**F**) CytoCatch archives POIs and resolves their abundance across time via an additional backfiller component to enforce forward-only recruitment of newly appearing POIs.

## 2 Results

### 2.1 Recorder of cytosolic protease activity

Proteases are dynamic and essential mediators of cellular physiology, regulating signal transduction and cell homeostasis^27^, with dysregulation implicated in diseases such as cancer^28^ and neurodegeneration^29^. Viruses exploit such machinery in the host cell, often encoding and expressing their own proteases, to facilitate replication, cell entry, and immune evasion^30^. Proteases have been identified as a major therapeutically targetable protein class^31^, and measuring their activity is essential for understanding their cellular functions and for evaluating therapeutic interventions^32,33^. We first set out to test whether protein tape recording of cytosolic protease activity could be achieved. To this end, we leveraged the Small Molecule-Assisted Shutoff (SMASh) sequence^34^, which comprises the hepatitis C virus (HCV) nonstructural protein 3 (NS3) protease (NS3pro) flanked by its cleavage site and a protein degradation sequence (the HCV NS4A degron)^35^. In the absence of asunaprevir (ASV, an NS3 protease inhibitor for treating HCV infection), the protease cleaves at its cleavage site and removes itself and the degron from the protein fused to the N-terminus of the SMASh sequence. Upon ASV treatment, the protease activity is inhibited, resulting in retention of the degron and subsequent degradation of the SMASh-fused protein.

We fused the SMASh sequence to the C-terminus of the V5-tagged CytoTape monomer^15^, resulting in a CytoSense system signal monomer, CytoSense-NS3pro, driven by the constitutive human ubiquitin C (*UbC*) promoter to maintain a stable level of signal monomer expression over time for sensing and reporting events in the cytosol (**Fig. 2A**). As the CytoTape assembly continuously elongates over time, we hypothesized that the NS3pro protease activity would be directly sensed by CytoSense-NS3pro and subsequently encoded as changes in this V5-tagged monomer density along the tape. Previous work has shown that 3 days of sustained ASV treatment in cultured neurons induced a substantial reduction in the abundance of the protein fused to the SMASh sequence^34^. Consistently, we observed a significant reduction of signal monomer density towards the ends of protein tapes under 3 days of ASV treatment in cultured neurons co-expressing CytoSense-NS3pro and CytoTape structural monomer, compared to the untreated control (**Fig. 2, B** to **E**).

**Fig. 2.**
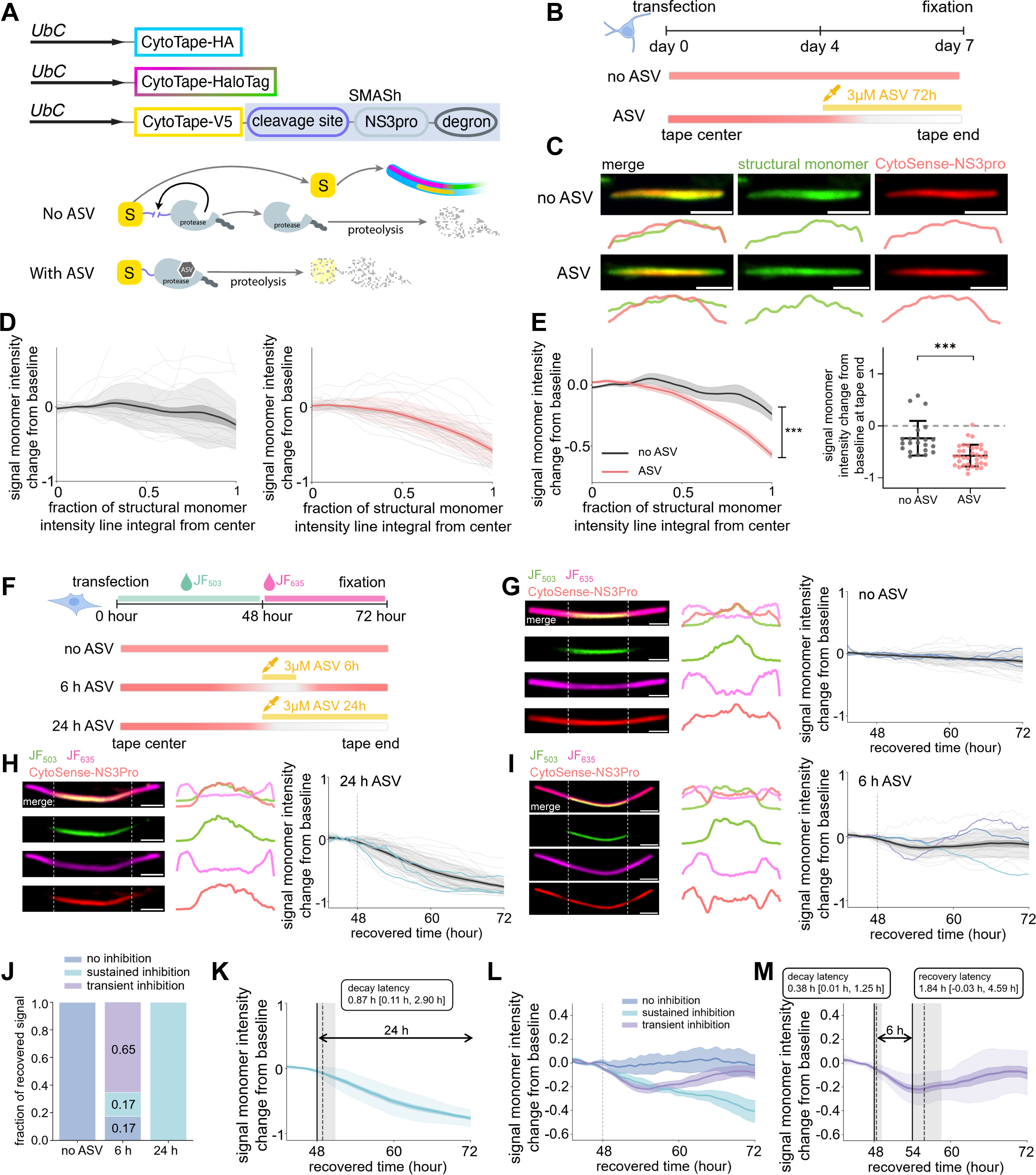
CytoSense enables temporally resolved recording of protease activity. (**A**) Schematic of the CytoSense-NS3pro system. The CytoSense-NS3pro monomer was constructed by fusing the SMASh sequence, comprising NS3pro flanked by its cleavage site and a constitutive degron, to the V5-tagged CytoTape monomer, driven by the constitutive *UbC* promoter. Protease activity cuts the cleavage site and removes the degron, modulating the stability of the signal monomer (yellow) population for durable incorporation into the elongating protein assembly. When protease activity decreases, fewer signal monomers are cleaved, resulting in degron retention and thus degradation of an increasing number of signal monomers before incorporation into the assembly. NS3pro, HCV nonstructural protein 3 protease; ASV, asunaprevir, a protease inhibitor of NS3pro; SMASh, Small Molecule-Assisted Shutoff sequence. (**B**) Top panel, experimental timeline for recording the protease activity of NS3pro in cultured neurons; bottom panel, expected CytoSense-NS3pro monomer density profile from tape center to tape end. (**C**) Representative confocal images of CytoSense-NS3pro tapes in the no-ASV control group (top) and after sustained 72 h ASV treatment (bottom) in cultured neurons. (**D**) Signal monomer intensity change from baseline for individual tapes and population average for no-ASV control (left panel, *n* = 22 tapes from 22 cells) and ASV-treated experiment (right panel, *n* = 35 tapes from 35 cells) groups. Throughout this figure unless specified: center line, mean; dark shaded boundary, s.e.m.; light shaded boundary, s.d. (**E**) Comparison between no-ASV control (gray) and ASV-treated experiment (red) groups on their signal monomer intensity change from baseline across time (left; shaded boundary, s.e.m.) and at the end of tapes (right; mean ± s.d.). ***P < 0.001; two-sided Mann-Whitney U test. (**F**) Top panel, experimental timeline for recording the protease activity of NS3pro in U2OS cells with HaloTag-based timestamp, co-transfected with CytoTape-vivo as the strucutral mononer; bottom panel, expected CytoSense-NS3pro monomer density profile from tape center to tape end. (**G**) Representative confocal images of CytoSense-NS3pro tapes (left), their fluorescence intensity profiles (middle), and population-level signal monomer intensity change from baseline across recovered time (right) in U2OS cells in the no-ASV control group. *n* = 27 tapes from 27 cells. Right panels in **G-I**: traces from three representative tapes in each group are highlighted in dark blue (for no inhibition), purple (for transient inhibition), or light blue (for sustained inhibition); traces from other individual tapes are in gray; vertical dashed line, onset time of ASV treatment. (**H**) Same as **G**, but for the 24 h ASV treatment group. *n* = 34 tapes from 34 cells. (**I**) Same as **G**, but for the 6 h ASV treatment group. *n* = 23 tapes from 23 cells. (**J**) Relative composition of temporal modes of recorded responses. Numbers in the bars, proportions. (**K**) Characterization of response latency for the 24 h ASV treatment group in **H**. Vertical solid line, onset time of ASV treatment; vertical dashed line and vertical shaded region, median and interquartile range (IQR) of signal decay onset time.The black box indicates the median and IQR of the latency between ASV treatment and signal decay onset. (**L**) Comparison of distinct temporal modes in the 6 h ASV treatment group in **I**. Dark blue, no inhibition, *n* = 4 tapes from 4 cells; purple, transient inhibition, *n* = 15 tapes from 15 cells; light blue, sustained inhibition, *n* = 4 tapes from 4 cells; center line, mean; shaded boundary, s.e.m.; vertical dashed line, onset time of ASV treatment. (**M**) Characterization of response latency for responses in the transient inhibition mode in **L**. Solid black lines, onset times of ASV treatment (48 h) and ASV withdrawal (54 h); vertical dashed lines and vertical shaded regions, medians and IQRs of response onset times for signal decay and recovery. Black boxes indicate the median and IQR of the latency between ASV treatment and signal decay onset and between ASV withdrawal and signal recovery onset, respectively. Scale bars throughout this figure, 5 μm.

We next expressed CytoSense-NS3pro in a proliferating human osteosarcoma cell line, U2OS, and characterized its temporal kinetics. We employed the HaloTag-based timestamp monomer from the CytoTape toolkit in the CytoSense system to provide the time axis along the tape via switching of distinct HaloTag ligand dyes at defined time points (**Fig. 2F**)^15^. Comparing among XRI- and CytoTape-derived monomer designs, we found that the CytoTape and CytoTape-vivo monomer designs produced thin, long, and flexible linear assemblies (referred to as “tapes” throughout this paper), with CytoTape-vivo having the most uniform morphological metrics across individual U2OS cells (**Fig. S1**). We therefore applied CytoTape-vivo for implementing CytoSense-NS3pro recording of NS3pro protease activity in U2OS cells.

When a 24-hour-long sustained ASV treatment was initiated concurrently with the HaloTag-based timestamp (via switching of HaloTag ligand dyes in the cell culture media) at 48 h after transfection, the onset of the decay of signal monomer density was closely aligned with the timestamp while progressing to a strong, sustained signal reduction along the tapes (**Fig. 2H**). Across the cell population, the median onset time recovered by HaloTag timestamp across individual tapes was 48.87 h after transfection, corresponding to a median onset latency of 0.87 h relative to the onset of ASV treatment (IQR, 0.11–2.90 h; **Fig. 2K**). These results indicate that the recorder is capable of capturing an immediate decrease in protease activity with at most hour-scale uncertainty of temporal latency after the onset of ASV treatment.

We next applied a transient 6-h ASV treatment in U2OS cells, with ASV being thoroughly washed out to minimize its further inhibition of protease activity after ASV withdrawal (**Fig. 2I**). In the majority of cells, CytoSense-NS3pro reported a clear, immediate “dark band” of signal monomer density along the tape that was temporally aligned with the 6-h treatment period, followed by recovery of signal monomer density after ASV withdrawal (**Fig. 2, J** and **L**). Interestingly, the 6-h ASV treatment group also contained cells that showed either no detectable signal reduction upon ASV treatment or sustained signal reduction without full recovery to the pre-treatment level after ASV removal, unlike the 24-h ASV treatment group where all cells exhibited the same sustained signal reduction behavior (**Fig. 2, J** and **L**). This unique mosaic pattern in signal recovery in the 6-h treatment group highlights cell-to-cell variation in responsiveness upon transient ASV treatment, potentially due to the heterogeneity in intracellular ASV clearance, protein turnover, or cell-specific proteostasis, as previously reported^36,37^. Among cells exhibiting transient signal reduction, the recorded signal exhibited an immediate decrease upon the onset of ASV treatment, consistent with the previous observation in the 24-h treatment group (**Fig. 2M**). We also quantified the recovery onset, defined as the time at which signal intensity began to increase, reflecting the recovery of signal monomer stability and incorporation into the tape. The median recovery latency, which is the interval between ASV withdrawal and recovery onset, was 1.84 h (IQR, −0.03 to 4.59 h) (**Fig. 2M**).

Together, these results demonstrated that the CytoSense system can utilize the signal monomer of the protein tape recorder as a direct activity sensor to record cytosolic protease activity with at most hour-scale median latency. More broadly, they established the adaptability of protein tape recorders for tracking temporal changes in signal monomer abundance when its degradation prior to tape incorporation is dynamically coupled to cellular activity of interest.

### 2.2 Recorder of mTORC1–S6K signaling activity

We next sought to determine whether the CytoSense system is generalizable to record endogenous cytosolic signaling dynamics, exploiting signaling activity-responsive degrons. We focused on the mTORC1 signaling axis, a master pathway that governs many essential aspects of cellular processes, such as protein synthesis^38^, metabolism^39^, and cell growth, proliferation, and survival^40,41^. Dysregulation of mTORC1 activity is implicated in a wide range of physiological conditions and diseases, whereas pharmacological inhibition of mTORC1 signaling has shown potential in contexts such as cancer intervention and anti-aging^42–44^. One of the major downstream effectors of mTORC1 signaling is S6 kinase (S6K) activation, which in turn activates ribosomal protein S6 and simultaneously phosphorylates translational repressor PDCD4 at its phosphodegron^45,46^. Phosphorylated PDCD4 is subsequently recognized by the ubiquitin-proteasome system and undergoes rapid proteasomal degradation, thereby lifting the brake on translation initiation in the presence of active cellular mTORC1 signaling. Conversely, inhibition of mTORC1 signaling suppresses S6K activity, maintains the PDCD4 phosphodegron in a hypophosphorylated state, and thus protects PDCD4 from degradation. Therefore, the phosphodegron of PDCD4 has been established as a dynamic readout of mTORC1–S6K signaling activity, as demonstrated by FP fusions of this phosphodegron as inverse activity probes^47,48^.

We reasoned that fusing the PDCD4 phosphodegron motif to CytoTape would yield a signal monomer whose cytosolic stability, and subsequent incorporation onto the protein tape, is modulated by the activity of mTORC1–S6K signaling in an inverse manner.

Under high mTORC1–S6K activity, enhanced phosphorylation of PDCD4 phosphodegron would reduce signal monomer stability and tape incorporation, whereas low mTORC1–S6K activity would stabilize the signal monomer and increase its accumulation along the protein tape. In this way, endogenous cytosolic signaling dynamics would be transformed into stable spatial records embedded along intracellular protein tapes (**Fig. 3A**).

**Fig. 3.**
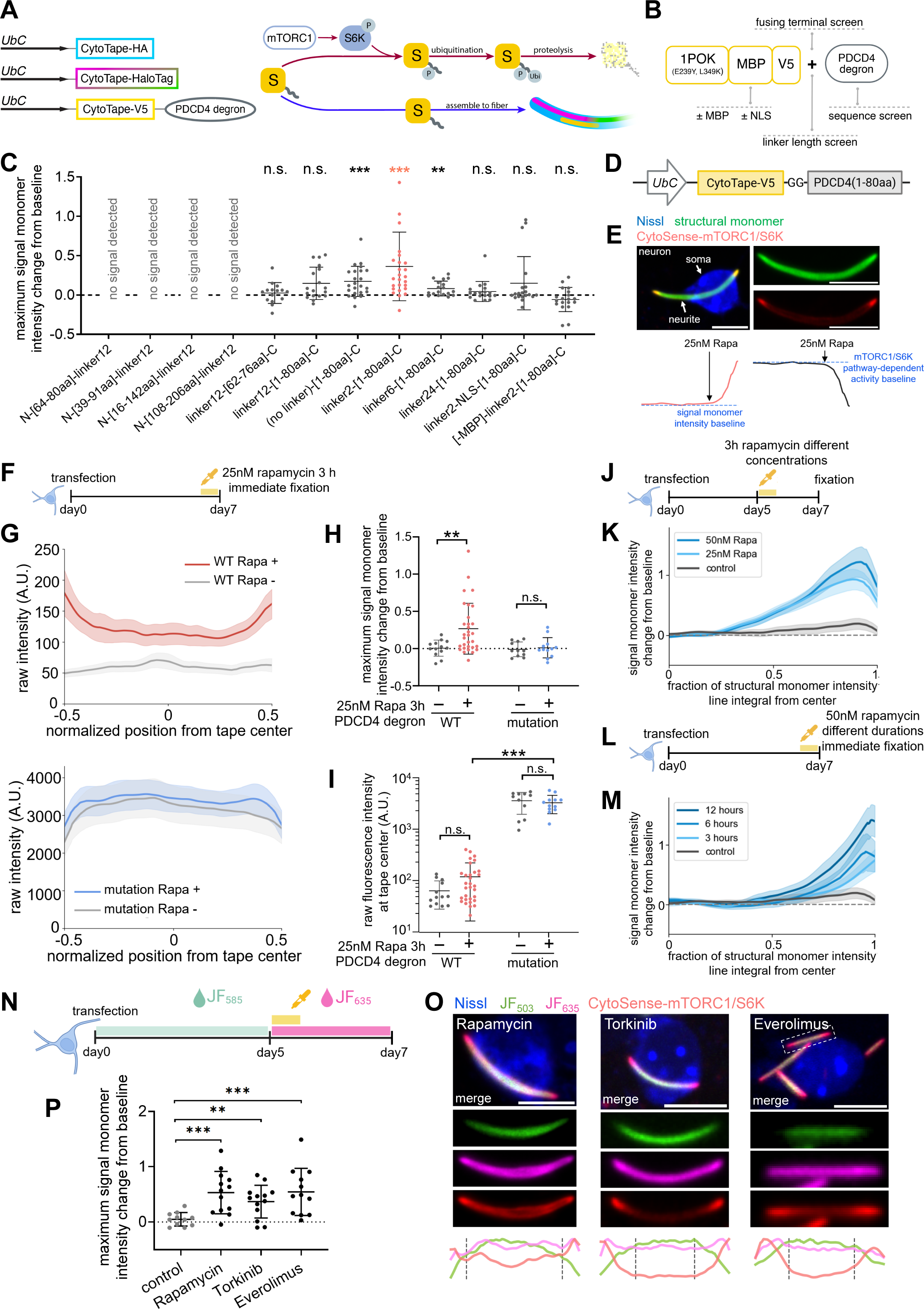
CytoSense enables temporally resolved recording of mTORC1–S6K signaling pathway activity. (**A**) Design of the CytoSense-mTORC1/S6K recorder system. The CytoSense-mTORC1/S6K monomer (yellow) was constructed by fusing a PDCD4-derived phosphodegron to the V5-tagged CytoTape monomer, driven by the constitutive *UbC* promoter. mTORC1–S6K signaling activity modulates phosphorylation-dependent degradation of signal monomers before their incorporation into the elongating protein assembly. Once incorporated, signal monomers are protected from degradation, providing a durable temporal record of signaling activity. (**B**) Design parameters tested for CytoSense-mTORC1/S6K engineering, including degron sequences, fusion terminus, linker length, and monomer architectural modifications such as MBP or NLS addition or deletion. (**C**) Comparison of the maximum signal monomer intensity change from baseline after rapamycin (Rapa; an mTORC1 inhibitor) treatment (mean ± s.d.) across distinct construct designs in cultured neurons. The design candidate highlighted in red is the selected design for CytoSense-mTORC1/S6K (also illustrated in **D**). *n* = 15 (linker12-[62–76aa]-C), 18 (linker12-[1–80aa]-C), 23 ((no linker)-[1–80aa]-C), 26 (linker2-[1–80aa]-C), 18 (linker6-[1–80aa]-C), 18 (linker24-[1–80aa]-C), 20 (linker2-NLS-[1–80aa]-C), and 19 ([-MBP]-linker2-[1–80aa]-C) tapes from the same numbers of cells, respectively. n.s., not significant; **, P < 0.01; ***, P < 0.001; one-sample Wilcoxon signed-rank tests against the baseline intensity level (“0.0”) with Holm correction for multiple comparisons. (**D**) Construct design of the CytoSense-mTORC1/S6K signal monomer. (**E**) Top panel, representative confocal images of CytoSense-mTORC1/S6K from screening in **C**. Bottom panel, recorded signal monomer intensity change from baseline along the tape (red, left) and the corresponding recovered mTORC1–S6K signaling activity (black, right; by inverting the intensity profile from the left). Blue dashed lines, baseline for each corresponding condition. (**F**) Experimental timeline for CytoSense-mTORC1/S6K recording in cultured neurons for results shown in **G**. (**G**) Raw fluorescence intensity profiles along the tape for CytoSense-mTORC1/S6K (top; “WT”) and its degron-inactivated mutant (PDCD4 degron with S67A, S71A mutations) (bottom; “mutation”), with (top red, bottom blue) or without (gray) rapamycin treatment. Top red, *n* = 31 tapes from 31 cells; top gray, *n* = 13 tapes from 13 cells; bottom blue, *n* = 12 tapes from 12 cells; bottom gray, *n* = 11 tapes from 11 cells. Throughout this figure: center line, mean; shaded boundary, s.e.m. (**H**) Quantification of the maximum signal monomer intensity change from baseline for CytoSense-mTORC1/S6K or its degron-inactivated mutant with (+) or without (−) 25 nM rapamycin treatment for 3 h (mean ± s.d.). (**I**) Quantification of raw fluorescence intensity at the center of the tapes from **H**. (**J-M**) Signal monomer intensity change from baseline plotted across the fraction of the line integral of structural monomer intensity for CytoSense-mTORC1/S6K recording under rapamycin treatment with distinct concentrations (**K**), no-treatment control, gray, *n* = 11 tapes from 11 cells; 25 nM, light blue, *n* = 30 tapes from 30 cells; 50 nM, dark blue, *n* = 30 tapes from 30 cells, following the experimental timeline in **J**; or with distinct durations (**M**), no-treatment control, gray, the same control dataset from **K** was plotted again here for reference; 3 h, light blue, *n* = 27 tapes from 27 cells; 6 h, blue, *n* = 16 tapes from 16 cells; 12 h, dark blue, *n* = 30 tapes from 30 cells, following the experimental timeline in **L**. (**N**) Experimental timeline for CytoSense-mTORC1/S6K recording with HaloTag-based timestamps in cultured neurons. (**O**) Representative confocal images of CytoSense-mTORC1/S6K recording (top rows), and their fluorescence intensity profiles (bottom row, gray dashed lines indicate timestamp points), following 2 h of 25 nM rapamycin treatment (left), 1 μM torkinib treatment (middle), or 50 nM everolimus treatment (right), under the timeline shown in **N**. (**P**) Quantification of the maximum signal monomer intensity change from baseline recorded by CytoSense-mTORC1/S6K under each treatment (mean ± s.d.) in **O**. Control, *n* = 11 tapes from 11 cells; rapamycin-treated group, *n* = 13 tapes from 13 cells; torkinib-treated group, *n* = 13 tapes from 13 cells; everolimus-treated group, *n* = 12 tapes from 12 cells. Throughout this figure: n.s., not significant; **, P < 0.01; ***, P < 0.001; two-sided Mann-Whitney U test. Scale bars throughout this figure, 10 μm.

Although the core PDCD4 phosphodegron is a short N-terminal motif spanning amino acids 70-76 of the PDCD4 protein, surrounding residues are also known to contribute substantially to effective degron function as they are involved in phosphorylation and substrate recognition in the proteolysis process^45^. We therefore screened distinct truncations of the N-terminal region of mouse PDCD4 in cultured neurons, to determine which configuration would yield the most robust responsiveness to mTORC1–S6K activity when fused to either the N- or C-terminus of the CytoTape monomer^49^ (**Fig. 3B**). To perturb the mTORC1 signaling activity for this screening, we used rapamycin, a well-characterized FDA-approved allosteric inhibitor of mTORC1 with broad relevance to immunosuppression, cancer therapy, and anti-aging-related interventions^50–52^. After treatment of 25 nM rapamycin for 3 h in cultured neurons at 7 days post-transfection, we immediately fixed cells for immunofluorescence imaging of CytoTape monomers. We found that fusion of PDCD4 amino acids 1-80 to the C terminus of the CytoTape monomer produced the strongest signal response among the tested signal monomer designs (**Fig. 3C** and **Fig. S2**). Interestingly, all designs with N-terminal fusions of PDCD4 truncations to the CytoTape monomer showed no detectable presence of signal monomer in the cells. We next tested whether the choice of protein domain linker between the CytoTape monomer and PDCD4(1-80 aa) might alter the efficiency of degron phosphorylation and recognition by cellular degradation machinery. Through systematic linker screening ranging from 0 (no linker) to 24 amino acids (aa) in linker length, we identified that constructs with no linker and with a 2- or 6-aa linker exhibit significant signal responses to rapamycin treatment (**Fig. 3C** and **Fig. S2**). The construct with the 2-aa linker (Gly-Gly) had the largest signal response, and we therefore refer to this signal monomer design (with PDCD4(1-80 aa) and 2 aa linker) as CytoSense-mTORC1/S6K throughout this paper (**Fig. 3, D** and **E**).

To validate that the responsiveness of CytoSense-mTORC1/S6K arises from the phosphodegron mechanism, we mutated two key phosphorylation target sites for S6K within the PDCD4 phosphodegron region (S67A and S71A) in CytoSense-mTORC1/S6K, as these mutations have been reported to impair the phosphodegron function (**Fig. 3, F** and **G**)^45^. The mutations completely abolished the recorder’s response to rapamycin, indicating that CytoSense-mTORC1/S6K is sensitive to mTORC1–S6K activity via the phosphodegron mechanism (**Fig. 3H** and **Fig. S3**).

The recorded pre-stimulation baseline intensity of the signal monomer with the two mutations was also significantly elevated compared to that of unmutated CytoSense-mTORC1/S6K, indicating that the unmutated phosphodegron in CytoSense-mTORC1/S6K remains sensitive and responsive to basal, physiological levels of mTORC1–S6K activity in neurons (**Fig. 3, G** and **I**). Importantly, these results support that degron-fused signal monomers that had already been incorporated along the tape remained stable across time, potentially due to the tape assembly insulating monomers from being accessed by the protein degradation machinery. Such stability of archived monomers provides the foundation for durable encoding of temporal histories along the protein assembly.

To further characterize CytoSense-mTORC1/S6K, we treated neurons with either 25 nM or 50 nM rapamycin for 3 h at 5 days post-transfection and fixed the neurons at 7 days post-transfection. The recorded inhibitory signal exhibited a dose-dependent increase with rapamycin concentration (**Fig. 3, J** and **K**). In addition, when cells were treated with 25 nM rapamycin for 3, 6, or 12 h at 7 days post-transfection, followed immediately by fixation at the end of each treatment period, the recorded inhibitory signals increased progressively with increasing treatment durations (**Fig. 3, L** and **M**). These results support CytoSense-mTORC1/S6K as an analog recorder of mTORC1–S6K activity level. To examine whether CytoSense-mTORC1/S6K broadly captures pharmacological perturbations of the mTORC1–S6K pathway and reliably reports the timing of such perturbations, we next treated neurons with rapamycin or one of the two other well-established mTOR-associated inhibitors, everolimus (also named RAD001, a rapalog-type allosteric mTORC1 inhibitor FDA-approved for immunosuppression and anti-cancer treatment^53^) and torkinib (also named PP242, a potent ATP-competitive mTOR inhibitor^54^), together with simultaneous HaloTag-based timestamping at the onset time of each pharmacological perturbation (**Fig. 3N**). As expected, all inhibitors induced an immediate signal response of CytoSense-mTORC1/S6K on the tape, with the response onset time aligned with the HaloTag-based timestamp (**Fig. 3O**) and peak signal response significantly larger than that in the no-perturbation control group (**Fig. 3P**). These observations support the temporal fidelity of CytoSense-mTORC1/S6K in capturing diverse pharmacological perturbations of mTORC1–S6K signaling.

Collectively, these results establish CytoSense-mTORC1/S6K as a genetically encoded, temporally continuous, longitudinally intensity-resolved recorder of mTORC1–S6K signaling dynamics, enabled by efficient activity-dependent modulation of signal monomer stability via a PDCD4-derived phosphodegron.

### 2.3 Validation and analysis of recorded mTORC1–S6K signaling dynamics

To validate the temporal recording fidelity of CytoSense-mTORC1/S6K under pharmacological perturbation and physiological conditions, we performed a time-series sampling experiment in neurons under CytoSense-mTORC1/S6K recording, with anti-pS6 (Ser235/236) immunostaining as a ground-truth readout for mTORC1–S6K pathway activity in the same neurons at each temporal snapshot (**Fig. 4A**). As a steric inhibitor of mTORC1, rapamycin has been reported to exhibit slow intracellular clearance after transient treatment in various types of cells^55^. Consistent with this, anti-pS6 staining across transient (25 nM, 2 h) rapamycin-treated groups revealed sustained inhibition of the mTORC1–S6K pathway in most neurons, beginning immediately after rapamycin treatment and persisting throughout the 3-day post-treatment experiment window (**Fig. 4D** and **Fig. S4**). Particularly, a subset of neurons showed partial recovery of pathway activity towards the end of the recording window, indicating heterogeneity in the treatment recovery progression across neuron populations. Similarly, the signal recorded by CytoSense-mTORC1/S6K also showed an increase in the recorded inhibitory signal right after rapamycin exposure, followed by sustained signal elevation throughout the recording window in most tapes (**Fig. 4, B** and **C**). Towards the end of the 3-day recording window, a small subset of tapes showed reduced inhibitory signal, consistent with the observation in anti-pS6 staining (**Fig. 4, B** and **C**). In the no-rapamycin control group, the anti-pS6 staining fluorescence intensity remained high across the population (**Fig. 4D** and **Fig. S4**) and the recorded CytoSense-mTORC1/S6K signals remained relatively stable throughout the 8 sampling points (**Fig. 4C** and **Fig. S5**), indicating a steady basal mTORC1–S6K activity without pharmacological inhibition.

**Fig. 4.**
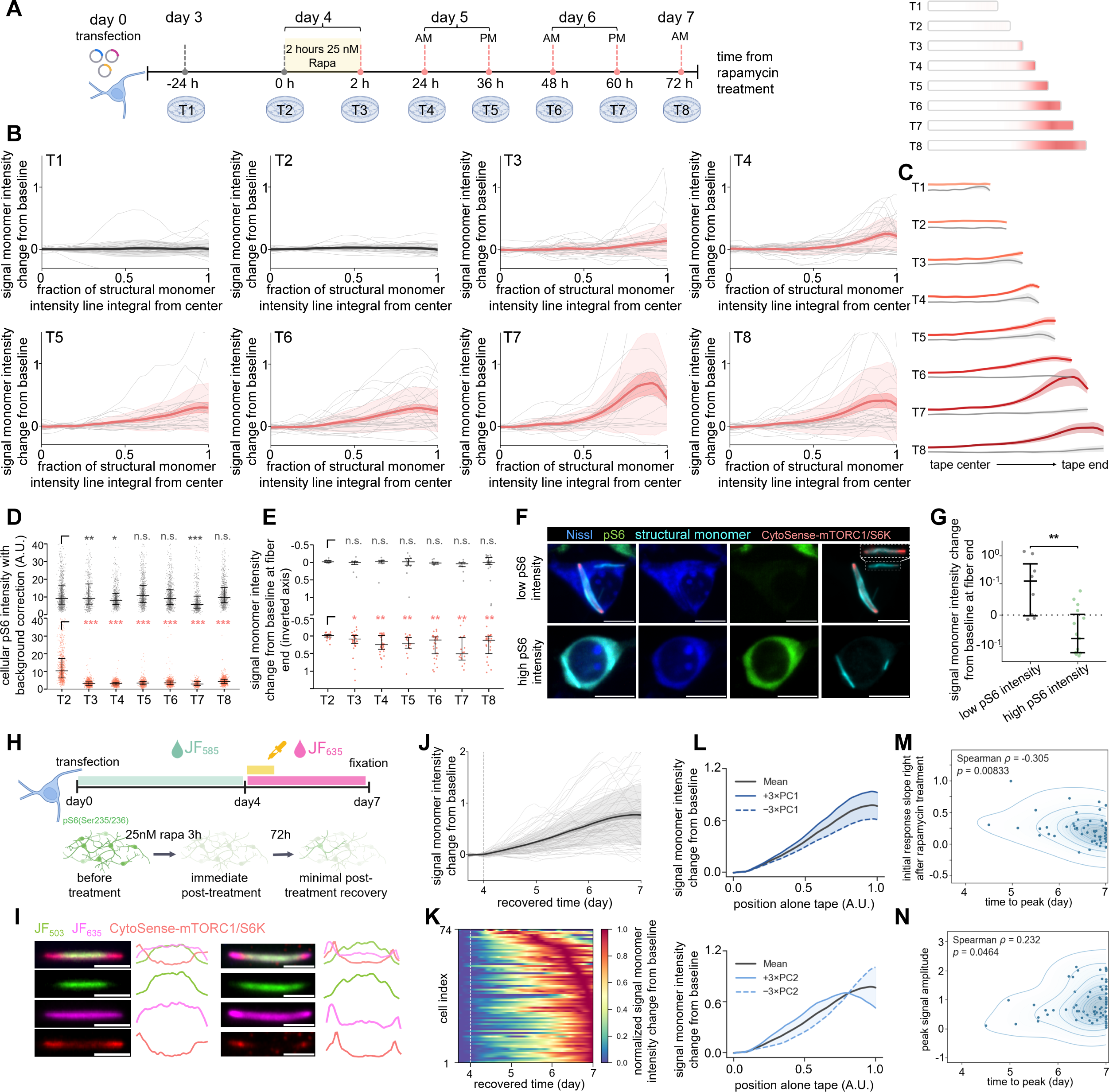
CytoSense recapitulates ground-truth immunostaining, provides physiological sensitivity, and resolves temporal features. (**A**) Experimental timeline for CytoSense-mTORC1/S6K recording and time-series sampling of mTORC1–S6K signaling pathway activity in cultured neurons. Neurons expressing the CytoSense-mTORC1/S6K system were transiently treated with rapamycin (25 nM, 2 h) at day 4 post-transfection and fixed at distinct time points from day 3 to day 7 (−24 h to 72 h relative to the onset of rapamycin treatment; labeled as T1-T8). The expected temporal progression of recorded signals along the tape at T1-T8 time points is illustrated on the right. (**B**) Signal monomer intensity change from baseline along individual tapes and population average for CytoSense-mTORC1/S6K recording across the 8 sampled time points (T1 and T2, gray, before stimulation; T3-T8, red, after stimulation). Throughout this figure unless specified: center line, mean; dark shaded boundary, s.e.m.; light shaded boundary, s.d.; gray thin lines, data from individual tapes. *n* = 33, 21, 32, 31, 20, 28, 20, 27 tapes from the same numbers of cells at T1-T8, respectively. (**C**) Signal monomer intensity change from baseline across T1-T8 (from top to bottom) for no rapamycin control (gray) and rapamycin-treated (gradient red, data adapted from **B**) groups. Center line, mean; shaded boundary, s.e.m. (**D**, **E**) Quantification of cellular anti-pS6 intensity from pS6 (Ser235/236) immunostaining (**D**) and signal monomer intensity change from baseline at the end of tapes (**E**) across the 8 sampled time points (T2-T8) in no rapamycin control (gray, top subplots) and rapamycin-treated (red, bottom subplots) groups of cultured neurons (median ± interquartile range). For **D**: control, *n* = 919, 510, 651, 703, 838, 627, 776 cells from 10, 10, 12, 8, 12, 12, 10 fields of view (FOVs) at T2–T8, respectively; rapamycin-treated, *n* = 858, 1134, 997, 530, 795, 312, 828 cells from 12, 16, 12, 12, 12, 10, 12 FOVs at T2–T8, respectively. For **E**: control, *n* = 11, 11, 10, 15, 11, 9, 25 tapes from the same numbers of cells at T2-T8, respectively; rapamycin-treated, *n* = 21, 32, 31, 20, 28, 20, 27 tapes from the same numbers of cells at T2-T8, respectively. n.s., not significant; *, P < 0.05; **, P < 0.01; ***, P < 0.001; two-sided Mann-Whitney U tests using the T2 group as control, with Holm correction for multiple comparisons. (**F**) Representative confocal images of CytoSense-mTORC1/S6K recorded neurons with low (top) or high (bottom) cellular anti-pS6 intensity. Scale bars, 10 μm. (**G**) Quantification of signal monomer intensity change from baseline at the end of tapes in neurons with low (gray, *n* = 8 tapes from 8 cells) or high (green, *n* = 15 tapes from 15 cells) cellular anti-pS6 intensity (mean ± s.d.). **, P < 0.01; two-sided Mann-Whitney U test. (**H**) Top panel, experimental timeline for CytoSense-mTORC1/S6K recording with HaloTag-based timestamps in cultured neurons. Bottom panel, expected cellular anti-pS6 intensity across time. (**I**) Representative confocal images of CytoSense-mTORC1/S6K tapes (left) and their corresponding fluorescence intensity profiles (right). Scale bar, 5 μm. (**J**) Signal monomer intensity changes from the baseline along the recovered time for the experiment described in **H**. *n* = 74 tapes from 74 cells. Dashed gray line, onset time of rapamycin treatment. (**K**) Heatmap of peak-normalized signal monomer intensity change from baseline recorded from individual tapes along the recovered time, from data in **J**. (**L**) The mean signal profile along the tapes from **J** together with trajectories reconstructed from ±3 principal component (PC) modes. Top plot, for PC1; bottom plot, for PC2. (**M**) Scatter plot with kernel density contours for the time to peak and the peak amplitude for results from **J**. Blue contours and background shading indicate kernel density estimate; darker shading, higher cell density. Spearman correlation coefficients (ρ) and two-sided p values are indicated in the panel. (**N**) Same plot as **M**, but for the time to peak and the signal slope within 12 h of rapamycin treatment onset.

Although each tape provides a continuous recording, to compare the recorded signal side-by-side with the temporally discrete, endpoint anti-pS6 immunostaining measurements within the same time window, we quantified the recorded signal at the ends of the tape (**Fig. 4, D** and **E**). Notably, although anti-pS6 and CytoSense-mTORC1/S6K report mTORC1–S6K activity through distinct mechanisms, we observed consistent temporal trends across both readouts. Specifically, in the control group without rapamycin treatment, both anti-pS6 staining and tape-end signals remained generally stable with modest fluctuation throughout the recording window. In the rapamycin-treated group, anti-pS6 staining and CytoSense-mTORC1/S6K both reported an immediate inhibition signal right after the 2-h rapamycin treatment, and this inhibition was sustained across the 3-day recording window relative to the pre-treatment baseline. Anti-pS6 immunostaining also revealed heterogeneous levels of physiological mTORC1–S6K activity across neurons in the control group without rapamycin treatment. To quantify whether CytoSense-mTORC1/S6K has sufficient sensitivity to detect such physiological fluctuations of mTORC1–S6K activity level, we further compared the anti-pS6 intensity and the CytoSense-mTORC1/S6K signal at the ends of tape in the same neurons in the control group at the final sampling time point. We found that neurons with minimal anti-pS6 fluorescence exhibited significantly higher tape-end inhibitory signal intensity than neurons with strong anti-pS6 fluorescence (**Fig. 4, F** and **G**). This correlation indicates that CytoSense-mTORC1/S6K can detect and distinguish mTORC1–S6K signaling activity levels with physiological sensitivity.

Because mTORC1 signaling regulates protein translation, we further investigated whether its inhibition affects the CytoSense-mTORC1/S6K recording that involves protein synthesis of monomers to support protein tape elongation. We quantified the elongation kinetics of tapes by characterizing their lengths across the experimental timeline. Tapes in rapamycin-treated and untreated groups showed comparable lengths at each sampled time point, with significantly longer tapes at the final sampled time point compared with the initial time point in both conditions (**Fig. S6, B** to **D**). These results indicate that mTORC1–S6K signaling dynamics induced by transient rapamycin treatment does not significantly alter tape elongation kinetics. Together, these data support CytoSense-mTORC1/S6K as a faithful recorder of mTORC1–S6K pathway dynamics in single cells across days under both pharmacological perturbation and physiological conditions.

We further characterized the temporal features of CytoSense-mTORC1/S6K recording across the 72 h window post-rapamycin treatment, with HaloTag-based timestamping to reconstruct the real-world time axis along the experimental timeline (**Fig. 4H**).

CytoSense-mTORC1/S6K revealed heterogeneous temporal trajectories of mTORC1–S6K signaling after transient (25 nM, 3 h) rapamycin treatment across individual neurons. Toward the end of the 72-h post-treatment window, some neurons exhibited partial recovery from the sustained signaling inhibition while others did not, with variable onset and recovery timing across neurons (**Fig. 4, I** to **K**). To dissect the major temporal features of single-cell variability, we performed principal component analysis (PCA) on the recorded trajectories (**Fig. 4L**). Principal component 1, which explained 85.09% of the variance, primarily captured differences in signal amplitude. Principal component 2, which explained 11.54% of the variance, captured differences in temporal dynamics, highlighting the degree of recovery from sustained signaling inhibition toward the end of the recording window. We then quantified PC1- and PC2-associated trajectory features, including peak signal amplitude, initial response slope right after rapamycin treatment, and time to peak (as a measure for the onset timing of signal recovery for neurons exhibiting such recovery). Correlation analyses showed that peak signal amplitude and initial response slope were weakly associated with time to peak (**Fig. 4, M** and **N**). Together, these results demonstrate that CytoSense-mTORC1/S6K records mTORC1–S6K signaling trajectories in a continuous and analog manner, enabling quantitative analysis of temporal dynamics across single cells.

### 2.4 Abundance-resolved temporal archive of proteins

The abundance of many cellular molecules both responds to upstream cellular activity and modulates downstream cellular processes, making its dynamics central for cell physiology and function. Tracking the dynamics of molecular abundance, beyond their activity, would provide a complementary and informative window into cellular processes. Here, we tested whether it is possible to physically capture cytosolic proteins and align them along elongating tapes to resolve changes in their abundance across time. To implement this concept, we fused a “catcher” domain to the CytoTape monomer to catch proteins of interest and archive them temporally along the tape through either target recognition interactions, such as nanobody-target recognition^56^, or engineered binding pairs, such as SpyCatcher-SpyTag with the 13-aa SpyTag motif fused to the protein of interest (POI)^57^. However, unlike signal monomer-based recording systems, which preserve temporal order by incorporation of signal monomers selectively to the ends of the elongating tape, we reasoned that the new protein archive concept may introduce the challenge of “backfilling”. Newly appearing proteins of interest might bind retroactively to unoccupied catcher domains on older segments of the tape, and thus disrupt the desired temporal order of archived molecules along the spatial axis of the tape. To address this, we introduced a constitutively expressed protein “backfiller”, which continuously and stably occupies unbound catcher domains on the tape and enforces forward-only molecular archiving along the tape.

Since *Fos* is a popular neuronal activity marker and *Fos*-dependent GFP reporter mouse lines are well established in neuroscience, we first tested this molecular archiving concept by capturing shEGFP expressed in primary cultured neurons from the TetTag transgenic mice^58^, a popular *Fos*-dependent shEGFP reporter mouse line. We fused the GFP nanobody^59^ to the C-terminus of the CytoTape monomer, driven by the constitutive *UbC* promoter, to continuously catch shEGFP. We found that this GFP nanobody-based CytoCatch system successfully captured and archived *Fos*-dependent shEGFP molecules on the tape (**Fig. S7, A** and **B**). We then added EGFP (Y66F; the non-fluorescent version^60^, referred to as EGFP(dark) hereafter), also driven by the constitutive *UbC* promoter, as a backfiller in the system. Upon forskolin (FSK) treatment to raise the neuronal *Fos* activity and thus increase the abundance of cellular shEGFP, this approach did not report a clear increase in shEGFP abundance relative to pre-FSK baseline (**Fig. S7, A** and **C**). Furthermore, these tapes exhibited a strong presence of the EGFP(dark) backfiller alongside only weak shEGFP fluorescence. We speculated that this lack of temporal order of archived molecules along the tape may be due to the reversible nature of the binding between the GFP nanobody and its targets^61^, a considerable difference in binding affinity between shEGFP and EGFP(dark), or a combination of both. This speculation was further supported by co-expression of *Fos*-dependent mEGFP with the GFP nanobody-based CytoCatch system via transfection in wild-type cultured neurons (**Fig. S7, A** and **D**), in which the tape predominantly archived mEGFP, while the EGFP(dark) backfiller distributed diffusively in the cell with negligible presence on the tape.

We then turned to the engineered covalent SpyCatcher-SpyTag binding pair, reasoning that its irreversible nature of binding reported in previous studies^57,62^ may overcome the issue observed with GFP nanobody and thus preserve the temporal order of archived molecules along the tape. We fused SpyCatcher, as the catcher domain, to the C terminus of the CytoTape monomer (referred to as the “CytoCatch-Spy monomer” hereafter), and fused its covalent binding motif, the 13-aa SpyTag, to a target POI, sfGFP (G67A, non-fluorescent version^63^, referred to as “sfGFP(dark)” hereafter). We referred to this CytoTape-SpyCatcher and SpyTag-POI system as CytoCatch-Spy hereafter. We next characterized the CytoCatch-Spy system in neurons by co-expressing *UbC* promoter-driven structural, timestamp, and CytoCatch-Spy monomers, together with *Fos* promoter-driven SpyTag-sfGFP(dark)-V5 as POI and the *UbC* promoter-driven SpyTag-sfGFP(dark)-FLAG as backfiller (**Fig. 5, A** and **B**). The CytoCatch-Spy system successfully captured the *Fos* promoter-dependent POI abundance change upon FSK treatment (**Fig. 5, E** to **G**), confirming that covalent capture, together with forward-only molecular archiving enforced by the presence of backfiller, provided an abundance-resolved temporal archive of proteins. In control experiments without forskolin stimulation, the archived POI abundance remained stable at a basal level along the tape (**Fig. 5, C** and **D**), suggesting that proteins archived on the tape remained stable without noticeable degradation, dissociation, or redistribution on the tape. Along the time axis recovered by HaloTag-based timestamping, the FSK-induced abundance increase of POI had multi-hour onset latencies across cells (with a median of 5.28 h; **Fig. 5H**).

**Fig. 5.**
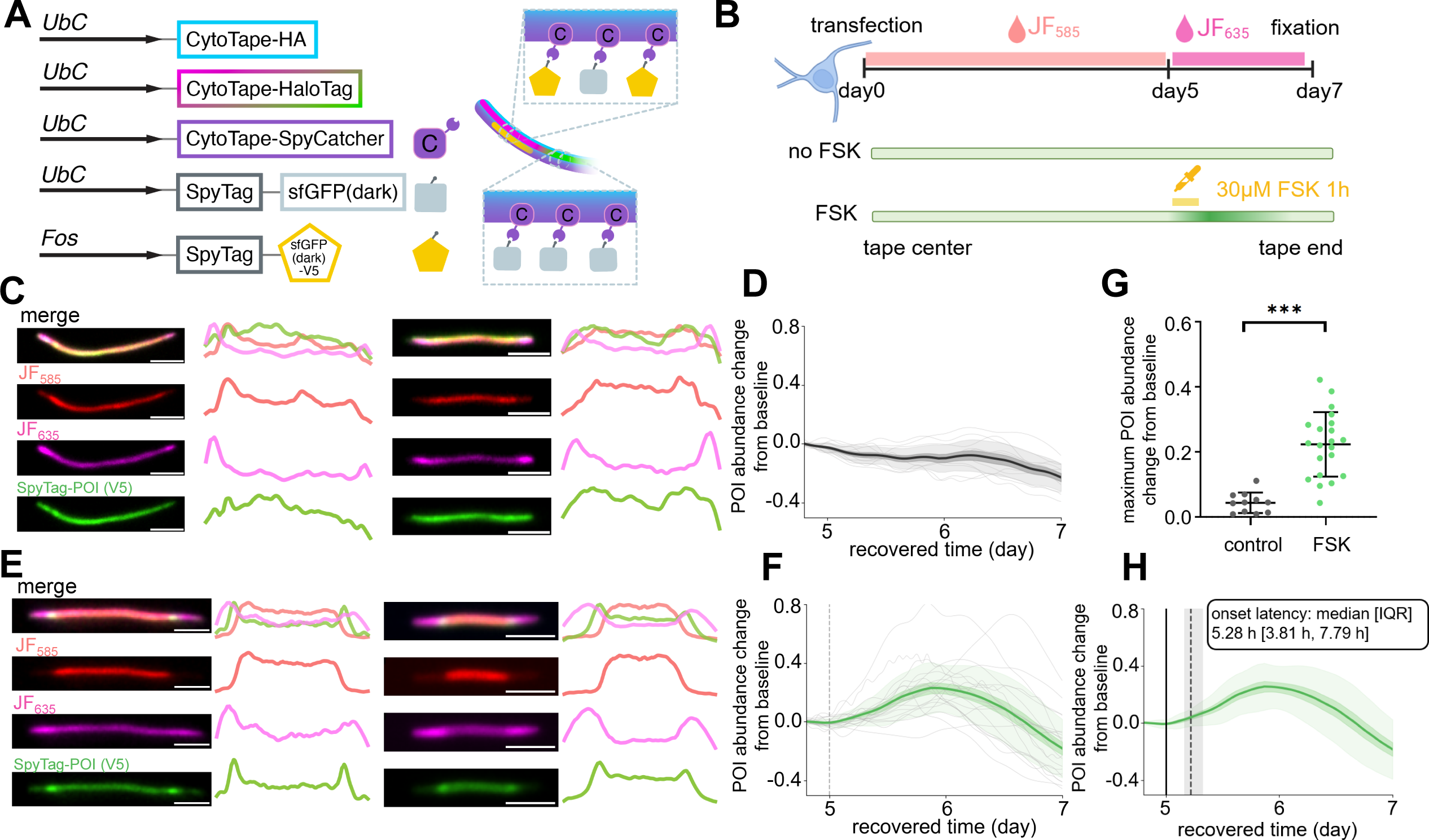
CytoCatch enables abundance-resolved, temporally resolved archives of cytosolic proteins. (**A**) Design of the CytoCatch-Spy system for an abundance-resolved temporal archive of cytosolic proteins. The CytoCatch-Spy monomer (CytoTape-SpyCatcher) was constructed by fusing SpyCatcher to the CytoTape monomer, driven by the constitutive *UbC* promoter. Proteins of interest (POIs) fused to the SpyTag (in this case, SpyTag-sfGFP(dark)-V5, driven by the activity-dependent *Fos* promoter) are covalently recruited onto the elongating tape, while a constitutively expressed backfiller, SpyTag-sfGFP(dark), enforces forward-only recruitment of newly appearing POIs along the tape. (**B**) Top panel, experimental timeline for CytoCatch-Spy archiving with HaloTag-based timestamps in cultured neurons; bottom panel, expected density profile of archived POIs from tape center to tape end. (**C**, **E**) Representative confocal images of archived POIs along CytoCatch-Spy tapes from the no-FSK control (**C**) and FSK-treated (**E**) groups (left), and their corresponding fluorescence profiles (right). Scale bar, 5 μm. (**D**, **F**) POI abundance change from baseline along the recovered time for no-FSK control (**D**) (gray, *n* = 11 tapes from 11 cells) and FSK-treated (**F**) (green, *n* = 21 tapes from 21 cells) groups. Throughout this figure unless specified: center line, mean; dark shaded boundary, s.e.m.; light shaded boundary, s.d.; gray thin lines, data from individual tapes; vertical dashed line, onset time of FSK treatment. (**G**) Quantification of the maximum POI abundance change from baseline for no-FSK control (gray, *n* = 11 tapes from 11 cells) and FSK-treated groups (green, *n* = 21 tapes from 21 cells) (mean ± s.d.). ***, P < 0.001; two-sided Mann-Whitney U test. (**H**) Characterization of response latency for the FSK-treated group in **F**. Solid black line, onset time of FSK treatment (day 5); vertical dashed line, median of response onset time for POI abundance change; vertical shaded region, interquartile range (IQR) of response onset time. The black box indicates the median and IQR of the latency between FSK treatment onset and response onset.

### 2.5 Temporal archive of molecular states and an internal clock

Beyond protein abundance, we next extended the CytoCatch system to analyze the temporal trajectory of molecular states of proteins, another key dimension of protein behavior in cells. To this end, we applied CytoCatch-Spy to track the chromophore maturation kinetics of fluorescent timers in the cytosol by physically capturing and archiving them along the tape for post hoc interrogation^21,64,65^. We co-transfected structural, timestamp, and CytoCatch monomers in U2OS cells without a backfiller, together with SpyTag-fused pSlow-FT^64^ as POI. pSlow-FT (referred to as “FT” hereafter) is a member of the fluorescent timer protein family, which naturally undergoes a spontaneous, gradual blue-to-red spectral transition after protein synthesis, with a previously reported red chromophore half-maturation time of ∼28 h in mammalian cells^64^ (**Fig. 6, A** and **B**). We imaged the cells 46 h after transfection and observed successful capture of FTs along the tape. Moreover, the tapes showed a robust color gradient from red to blue, extending from the tape center (older segments) towards tape ends (newer segments), indicating that this strategy preserves the temporal dynamics for FT chromophore maturation along the tapes (**Fig. 6C**). Considering the ∼28 h red chromophore half-maturation time of this FT, the robust color transition of FTs along the tape over our experimental timescale suggests the maturation process of FTs continued, rather than halted, after recruitment to the tape. The recorded temporal profiles of the red-to-blue ratio, an established measure of the maturation state of FTs^64,66^, showed conserved trends across cells and were consistent with previously reported maturation dynamics at the population level (**Fig. 6D**). We also observed tape-to-tape variation in the absolute red-to-blue ratio values. However, this variance was significantly larger between tapes from different cells than between tapes within the same cell (when there was more than one tape in a cell) (**Fig. 6E**). This suggests that tape-to-tape variation primarily arises from cell-to-cell heterogeneity in cellular conditions that may have influenced FT maturation. Together, these results demonstrated that CytoCatch-Spy can reliably preserve the temporal molecular states of FTs.

**Fig. 6.**
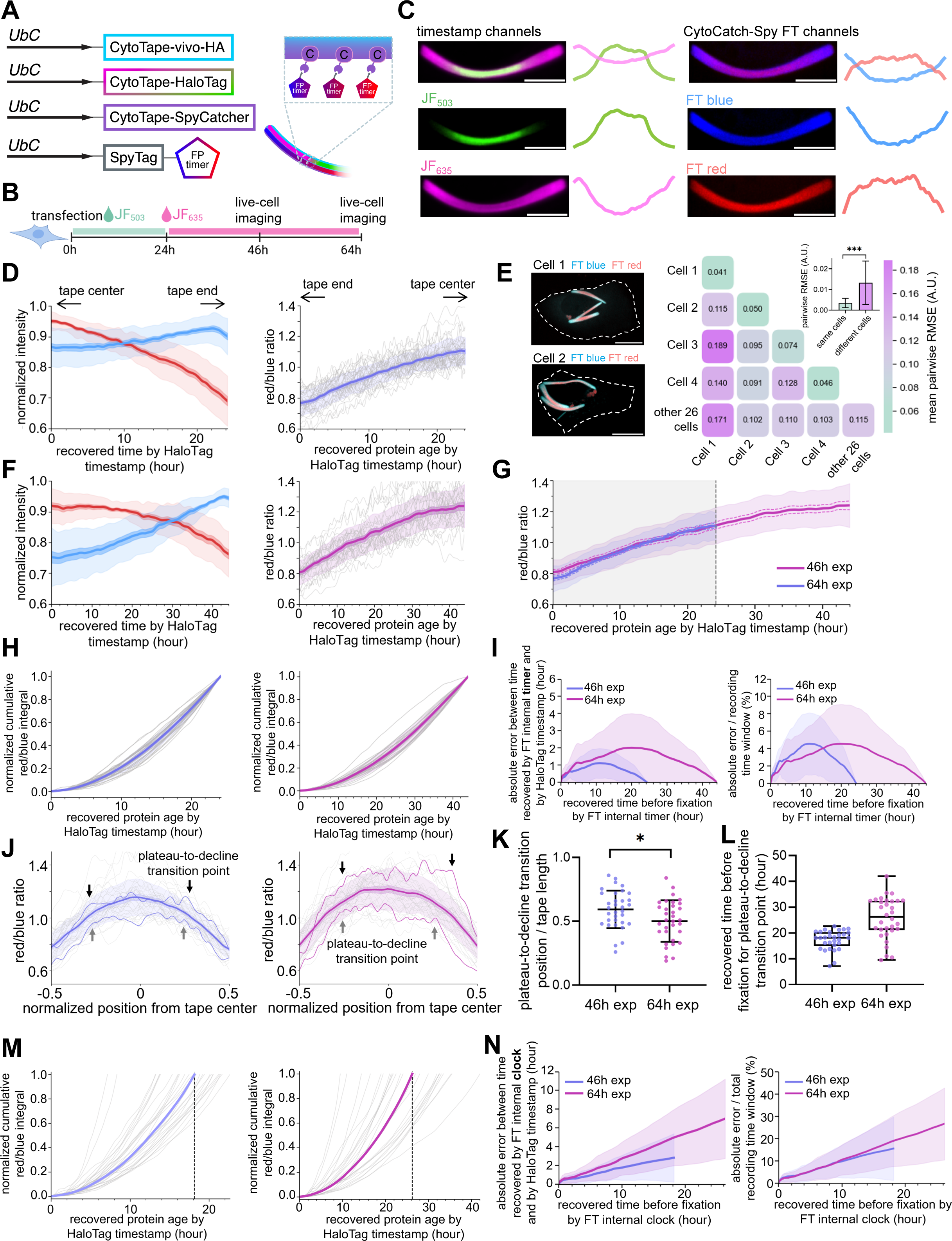
**CytoCatch tracks molecular state transitions of archived proteins and provides protein maturation-based internal clocks.** (**A**) Schematic of CytoCatch-Spy for temporal archiving of SpyTag-fused fluorescent timer (FT) proteins to track their blue-to-red chromophore maturation process across time. (**B**) Experimental timeline for CytoCatch-Spy temporal archiving with HaloTag-based timestamps in U2OS cells. Live-cell imaging was performed 46 h (for the 46 h experiment group) or 64 h (for the 64 h experiment group) post-transfection. (**C**) Representative confocal images of CytoCatch-Spy tapes with archived FTs in the timestamp (left) and FT (right) channels, alongside their corresponding fluorescence intensity profiles along the tape. Scale bar, 5 μm. (**D**, **F**) Left panels, normalized blue and red fluorescence intensity profiles of archived FTs along the recovered time for the 46 h (**D**) and 64 h (**F**) experiment groups. Throughout this figure unless specified: center line, mean; dark shaded boundary, s.e.m.; light shaded boundary, s.d.; gray thin lines, data from individual tapes. Right panels, red-to-blue ratio profiles along the recovered protein age by HaloTag timestamp for the 46 h (**D**, violet) and 64 h (**F**, purple) experiment groups. Throughout this figure: 46 h experiment group, violet, *n* = 30 tapes from 30 cells; 64 h experiment group, purple, *n* = 32 tapes from 32 cells. (**E**) Left panels, representative confocal images of cells containing multiple CytoCatch-Spy tapes. Dashed white line, cell boundary; scale bar, 10 μm. Heatmap on the right, quantification of mean pairwise RMSE between red-to-blue ratio profiles of the tapes within the same cells (Cells 1-4, *n* = 2, 3, 3, and 2 tapes per cell, respectively) and tapes from different cells (other cells, *n* = 26 tapes from 26 cells that had only 1 tape per cell). Top right bar plot, comparison of pairwise RMSE for tapes within the same cells (*n* = 8 pairs from Cells 1-4) versus tapes from different cells (*n* = 325 pairs) (mean ± s.d.). (**G**) Overlay of the red-to-blue ratio profiles from **D** and **F** along recovered protein age from HaloTag timestamp for the 46 h (violet) and 64 h (purple) experiment groups. Center line, mean; dashed line, s.e.m.; light shaded boundary around the center line, s.d.; light gray region with gray dashed line, recovered time window (24.2 h in duration) for the 46 h group. (**H**) Normalized cumulative integral of the red-to-blue ratio along recovered time for the 46 h (left, violet) and 64 h (right, purple) experiment groups. (**I**) Absolute error between time recovered by FT internal timer and by HaloTag timestamp (hour) for the 46 h (violet) and 64 h (purple) experiment groups calculated from **H**, using the mean profile of red-to-blue ratio as the transfer function of the internal timer. Left, absolute prediction error in hours. Right, absolute prediction error normalized to the total recording window (24.2 h and 44 h, respectively). Thick center line, mean; shaded boundary, s.d. (**J**) Red-to-blue ratio profiles along the tape for the 46 h (left, violet) and 64 h (right, purple) experiment groups. Thin violet and purple lines, data from individual representative tapes; black and gray arrows, plateau-to-decline transition points observed on the thin violet and purple lines. (**K**) Position of the plateau-to-decline transition point from the tape center relative to the half tape length for the 46 h (violet) and 64 h (purple) experiment groups (mean ± s.d.). (**L**) Time of the plateau-to-decline transition point, as recovered by HaloTag-based timestamp, for the 46 h (violet) and 64 h (purple) experiment groups (median ± interquartile range, with whiskers at extremes). (**M**) Normalized cumulative integral of the red-to-blue ratio from tape end to plateau-to-decline transition point as a function of the HaloTag-based recovered time for the 46 h (left) and 64 h (right) experiment groups. Gray thin lines, data from individual tapes; thick line, population median of 46 h (left) and 64 h (right) experiment groups, as the transfer function of the internal clock, see **Fig. S8D** for details of its calculation; vertical dashed line, median time of the plateau-to-decline transition point from **L**. (**N**) Absolute error between time recovered by FT internal clock and by HaloTag-based timestamp, as a function of time recovered by FT internal clock for the 46 h (violet) and 64 h (purple) experiment groups calculated from **M**. Left, absolute time error; right, absolute time error normalized by the total duration of the recovered time axis (18.15 h and 26.25 h, for 46 h and 64 h groups, respectively). Thick center line, mean; shaded boundary, s.d. Throughout this figure: *, P < 0.05; ***, P < 0.001; two-sided Mann-Whitney U test.

To evaluate temporal scalability, we replicated this experiment but instead imaged the cells 64 h after transfection, prolonging the duration of molecular capture (**Fig. 6F**). The red-to-blue ratio temporal profile from the prolonged experiment overlapped with that from the previous experiment (imaged 46 h after transfection) within their shared time window, then continued the same trend through the extended period, demonstrating the temporal scalability and reproducibility of this approach (**Fig. 6G**).

Inspired by the conserved temporal profiles of the red-to-blue ratio across cells and recording durations, we next examined whether they provide sufficient accuracy to serve as an internal indicator of time along the tape for real-world time axis recovery. Such a capability may empower future protein tape recording applications where external delivery of timestamps is challenging. By using the known start and end points of a recording window, defined in this case by the endpoint microscopy imaging time and a prior timestamp induced by the switching event of two HaloTag ligand dyes, we first asked whether FT could serve as an internal timer capable of recovering intermediate time points within the defined recording window. To establish a monotonic measure of time from the temporal profile of magnitude-normalized red-to-blue ratio, we calculated the cumulative integral of the temporal profile from tape center to tape end, and then averaged this cumulative integral across all tapes (**Fig. 6H**). The resulting averaged temporal profile was used as a transfer function between the cumulative integral across the tape (as a measure of the spatial axis along the tape^14^) and the time axis (recovered from HaloTag-based timestamping), to establish the internal timer to be applied along each tape. Within the defined time window, the mean absolute errors between the time axis recovered from the internal timer and that recovered from HaloTag timestamping were largest in the middle of the recording window and smallest at the beginning and the end, with error maxima of 1.16 h and 2.00 h for the 46-h and 64-h datasets, respectively, and remained below 1 h across most of the recovered time range (**Fig. 6I**). When normalized to the total duration of the time recovery window (defined as the time interval between HaloTag timestamping and endpoint imaging readout), the relative temporal accuracy was comparable between the two groups, with relative mean absolute errors of 4.81% and 4.54% for the 46-h (across the 24.2 h time recovery window) and 64-h (across the 44 h time recovery window) datasets, respectively (**Fig. 6I**). Together, these results demonstrate that archiving FT along protein assembly provides a reproducible internal timer with hour-scale accuracy across cells in a time recovery window with known start and end times.

Beyond its use as an internal timer, which requires a predefined time recovery window, we next asked whether FTs archived by CytoCatch-Spy could serve as a self-contained internal clock, enabling the recovery of time axis, and thus molecular age, without externally provided timestamps or other temporal references prior to the endpoint. We observed that most red-to-blue ratio profiles along the entire tape exhibited a characteristic two-phase profile, consisting of a stable plateau phase at the center of the tape followed by a declining phase toward the tape end, separated by a distinct plateau-to-decline transition point (**Fig. 6J** and **Fig. S8B**). By analyzing the positions of these plateau-to-decline transition points along the tape, we found that the transition point is experiment-duration dependent, as a longer experiment duration yielded transition points significantly closer to the tape ends (**Fig. 6K**). This two-phase profile closely mirrors previous reports on FT maturation kinetics, where the FP molecular population exhibits an ongoing maturation phase at an approximately linear rate over time before transitioning into a saturation phase that corresponds to the completion of maturation^64^. Thus, we reasoned that such a saturation of the red-to-blue ratio in the plateau region in the center (the earlier segment) of the tape may represent FT molecules that completed their maturation process, whereas the declining phase observed after the transition point represents newly synthesized FT molecules with the ongoing maturation process.

We therefore used the plateau-to-decline transition point in the red-to-blue ratio profile (referred to as the “transition point” hereafter) as an FT-derived timestamp. Using HaloTag-based time recovery, we converted the time to the transition point into the corresponding “transition time”, and the median transition times were 18.15 h and 26.25 h across individual cells from the 46-h and 64-h experiment groups, respectively (**Fig. 6L**). We then used the median transition time to define the time recovery window of the FT-based internal clock. To reconstruct molecular age from the FT-based internal clock without the information provided by HaloTag-based timestamping, we applied the same analysis pipeline used for the FT internal timer. Specifically, we calculated the cumulative integral of the normalized red-to-blue ratio profiles from the tape end to the transition point, and established the averaged cumulative integral profile across cells as the transfer function for time axis recovery (**Fig. S8, C** and **D**). We then evaluated its temporal accuracy by comparing the recovered time axis from the FT-based internal clock with that from the HaloTag-based timestamp in the same cells (**Fig. 6M**). The absolute prediction error increased with molecular age (defined as the elapsed time between the FT-derived time at a tape location and the experimental endpoint), with minimal error near the experimental endpoint and progressively increasing to several hours for time points further in the past (**Fig. 6N**). The lower temporal accuracy of FT-based internal clock compared with FT-based internal timer is expected, given that the time axis from the former was inferred from less known temporal reference information. Time errors likely also arise from cellular heterogeneity in red-to-blue ratio profiles, as characterized above (**Fig. 6E**).

Together, CytoCatch-Spy archives FT proteins along the tape and preserves their temporal progression of chromophore maturation, which further enables time axis recovery along tapes at hour-scale temporal accuracy across days, exploiting the archived FTs as an internal timer or an internal clock. The internal timer mode provides a higher temporal accuracy and requires a synchronized mark at the start of the recording window on each tape. The internal clock mode provides a lower temporal accuracy while eliminating the need for the synchronized “start” mark.

## 3 Discussion

Cytosolic processes play a central role in cellular physiology and function, and are highly dynamic over time^3^. Measuring cytosolic processes in single cells across time is critical for dissecting and understanding how cells sense, process, and respond to internal and external cues in healthy and diseased states. Molecular tape recording technologies have emerged as powerful platforms for scalable, time-resolved recording of cellular events^12^. However, existing approaches remain largely constrained by transcription-based recording mechanisms, in which transcriptional regulatory elements in the cell nucleus are used to sense cellular events, thus limiting their capability to capture cytosolic events, such as signaling pathways that are not closely coupled with transcription programs. Moreover, one often needs to examine the cytosolic molecules themselves across time, such as their molecular abundance and state changes, in addition to analyzing cellular activities. In this work, we introduced CyTRACE, a novel protein tape recording framework that utilizes cytosol-localized recording substrate blocks as direct activity sensors and cellular molecule catchers. Building upon this framework, we demonstrated two highly modular, genetically encoded systems:

CytoSense and CytoCatch, for temporally continuous and longitudinally intensity-resolved recording of cytosolic activities and physical archiving of cellular molecules, with physiological sensitivity, spatiotemporal resolution, and scalability.

Under the CyTRACE framework, we developed CytoSense, a system that transduces cytosolic activity into signal monomer stability via degron-mediated protein degradation, to be recorded as longitudinal changes in signal monomer density along the protein assembly for post hoc time-resolved readout. In this study, we demonstrated two representative implementations: CytoSense-mTORC1/S6K, for recording cellular mTORC1–S6K pathway-dependent activity, and CytoSense-NS3pro, for recording the HCV NS3 protease activity. Beyond these examples, CytoSense is modular by design, potentially enabling generalization to record a broader range of cellular and enzymatic activities, by fusing distinct cellular regulatory or enzymatic substrate domains to signal monomers. For example, CytoSense can be adapted to record cytosolic processes that can be coupled to degron-mediated regulatory mechanisms^67–70^. CytoSense may also be programmed to record a broader range of protease activity-dependent events for studies on cell physiology, disease, and therapeutic intervention^27^, such as those mediated by endogenous proteases or viral proteases expressed in infected host cells , by employing a signal monomer comprising simply a protease cleavage sequence flanked by the tape monomer and a constitutive degron sequence.

Such direct coupling of cellular activity to signal monomer stabilization and degradation may theoretically reduce the temporal delays inherent to transcription factor activation and transcriptional initiation in conventional molecule recording approaches. In this study, CytoSense-NS3pro exhibited sub-hour-scale response latency following NS3pro inhibitor treatment in U2OS cells when aligned to HaloTag-based timestamps. Of note, such latency characterization is inherently constrained by the temporal accuracy of the HaloTag timestamp-recovered time axis^15^. Similarly, CytoSense-mTORC1/S6K faithfully recorded sustained mTORC1 inhibition across days in the experiment window, which was aligned with the well-established mTORC1 activity marker pS6, following rapamycin treatment in neurons. Quantifying the exact temporal fidelity enhancement by cytosolic recording over transcriptional recording requires evaluation on a case-by-case, pathway-specific basis through side-by-side comparisons. Such comparisons are not straightforward, however, because the relationship between cytosolic activity dynamics and transcriptional outputs may vary substantially across pathways. Many biological events lack well-defined or direct transcriptional readouts, while others engage complex transcriptional programs with distinct delays, nonlinearities, and signal integration properties. As a result, a meaningful assessment of temporal fidelity would require carefully selected biological contexts and pathway-specific benchmarking frameworks, which would be an important direction for future investigation. Here we reason that the temporal fidelity of CytoSense is governed primarily by the kinetics of three processes: constitutive production of signal monomers, degron-mediated proteolysis, and incorporation of signal monomers into the growing protein tape. Future efforts will therefore benefit from systematic optimization and then quantitative characterization of these kinetic processes.

Under the CyTRACE framework, we developed CytoCatch-Spy, a molecular archive system that captures cytosolic proteins and aligns them along the elongating protein assembly, enabling scalable post hoc interrogation of their abundance and molecular states across time. In this study, CytoCatch-Spy captured protein abundance dynamics and resolved the temporal dynamics of protein maturation. In the latter case, a molecular clock was established with median temporal errors as low as 1.68 h for the 18.15-h duration and 3.51 h for the 26.25-h duration, via fluorescent timer proteins captured along the tape. This internal clock concept may potentially empower future protein tape recordings in applications where external delivery of timestamps is challenging.

The programmability of the catcher module makes this strategy readily adaptable to other protein targets, while also providing a potential path toward multiplexed cytosolic recording by capturing multiple molecular species simultaneously within single cells for scalable post hoc analysis. One approach would be to introduce orthogonal covalent binding systems, alongside SpyTag/SpyCatcher, enabling distinct protein species to be recruited through separate capture chemistries^71^. Alternatively, if the final readout is performed on the captured proteins themselves by methods such as immunostaining, it may also be feasible to fuse the same SpyTag to multiple proteins of interest and distinguish them after fixation using protein-specific antibodies. Future work may further exploit the accessibility of cellular molecules archived along the linear protein assembly to exogenous reagents, for retrospective interrogation of molecular properties across their molecular age and temporal history. Relatedly, a recent innovation archived transcriptome-wide mRNAs into vault particles at a chemically defined time point for post hoc analysis in cell-aggregated populations^72^. CytoCatch leverages the growing protein assembly as a novel molecular storage substrate that continuously archives target proteins for post hoc analysis across time. CytoCatch may further enable post hoc retrieval of archived molecules through controlled cleavage or dissociation from the protein assembly, for example by introducing a cleavage site or release module that can be activated chemically, enzymatically, optically, or by other defined physical or chemical stimuli.

Future extensions of CytoCatch-Spy for molecular state recording would also need to consider whether molecular archiving alters the ongoing molecular state transitions. Once recruited to the tape, a protein may not experience the exact same biochemical environment as its freely diffusing cytosolic counterpart^73^. In this study, the fluorescent timer was able to continue undergoing chromophore maturation after being archived, with kinetics similar to those reported in literature (a half-maturation time of tens of hours)^64^, suggesting that the incorporation of captured molecules onto the tape provides sufficient environmental contexts for the required chemistry. However, some state transitions or post-translational modifications may require the protein to have full access to other molecules or biochemical environments that archiving on the tape can no longer provide. In these cases, the transient molecular state present at the time of tape capture may be preserved until fixation, which could be exploited to enable recording of the temporal snapshots of their molecular states.

In summary, this study establishes CyTRACE as a novel molecular recording framework that transforms intracellular protein assemblies from transcriptional activity recorders into programmable archives of cytosolic activities and cellular molecules. Future studies of cellular processes would benefit from extending CyTRACE to broader biological contexts, including diverse cell types and animal models in both healthy and diseased states, as well as enabling multiplexed, longitudinal profiling of cytosolic activities and cellular molecules on the same tape. CyTRACE is also conceptually compatible with existing scalable molecular recording methodologies, such as the multiplexed transcriptional recorders of CytoTape^15^, the brain-wide recording framework GLOBE^74^, and multi-omics analyses, potentially enabling large-scale comprehensive spatiotemporal readout for in vitro and in vivo applications.

## 4 Methods

### Molecular cloning and human cell lines

Genes were synthesized by Twist Bioscience, and then inserted into mammalian expression vectors, pAAV-*UbC* and pAAV-*Fos*, from our previous work via Gibson assembly. See **Supplementary Table S1** for the sequence of the expressed genes and plasmid motifs. See **Supplementary Table S2** for constructs used in this study for different cell lines. Human cell lines used in this study include human bone osteosarcoma epithelial cells (referred to as “U2OS” throughout this paper; ATCC HTB-96).

### Animals and preparation of primary mouse hippocampal neuron cultures

All procedures in this work involving animals at the University of Michigan were conducted in accordance with the United States National Institutes of Health Guide for the Care and Use of Laboratory Animals, and were reviewed and approved by the University of Michigan Institutional Animal Care & Use Committee. Cultured neurons were prepared from neonatal (postnatal day 0 or 1) Swiss Webster mice (Taconic; both male and female mice were used) or B6.Cg-Tg(Fos-tTA,Fos-shEGFP)1Mmay/J (TetTag) mice (The Jackson Laboratory, stock no. 018306, used in **Fig. S7, B** and **C**), as previously described. Briefly, the brains of ice-anesthetized neonatal mice were dissected out, and the hippocampal tissue was further dissected from the brains in an ice-cold dissection buffer (1 mM Kynurenic acid, 10 mM MgCl_2_, 35 mM D-glucose in HBSS). The hippocampal tissue was subsequently treated with papain for 10 min at 37°C, followed by a 5-minute ovomucoid inhibition. The dissociated tissues were triturated, and a single-cell suspension was prepared using neuronal culture media (Minimum Essential Medium (MEM; Gibco) supplemented with 10 mM HEPES, 10% heat-inactivated fetal bovine serum (HI FBS; Corning), 2% B27 supplement (Gibco), 2 mM L-Glutamine, 1 µM transferrin, 25 mM D-glucose; pH adjusted to 7.4). Cells were plated at a density of 40,000 cells per 100 µL on Matrigel (Corning)-coated coverslips (Carolina) and then incubated in a humidified cell culture incubator at 37 °C with 5% CO_2_. After the cells had settled and attached to the coverslips, 1 mL of neuronal culture media was added.

When glial cells reached approximately 80% confluency (usually two days after plating), mL neuronal culture media supplemented with 4 µM AraC was added to the existing media in each well.

### CyTRACE expression in cultured neurons

Cultured neurons were transfected at 5-7 days *in vitro* (DIV 5-7) for structural monomer, timestamp monomer, and CytoSense or CytoCatch monomers. A commercial calcium phosphate transfection kit (Invitrogen) was used as previously described^14^, with the following transfection parameters. For experiments related to CytoSense-NS3pro (**Fig. 2, B** to **E**), each well of a 24-well glass-bottom plate received 100 ng of the structural monomer plasmid (*UbC*-CytoTape-HA), 20 ng of the timestamp monomer plasmid (*UbC*-CytoTape-HaloTag), 10 ng of the CytoSense-NS3pro monomer plasmid, and 1370 ng of pUC19 plasmid (a “dummy” plasmid without a mammalian open reading frame to maintain the optimal mass ratio between DNA and calcium phosphate for the formation of co-precipitates for transfection). For experiments related to CytoSense-mTORC1/S6K (**Fig. 3** and **Fig. 4**), each well of a 24-well glass-bottom plate received 100 ng of the structural monomer plasmid (*UbC*-CytoTape-HA), 20 ng of the timestamp monomer plasmid (*UbC*-CytoTape-HaloTag), 10 ng of the CytoSense-mTORC1/S6K monomer plasmid (or each candidate monomer plasmid during screening (**Fig. 3C**)), and 1370 ng of pUC19 plasmid. For experiments related to the GFP nanobody-based CytoCatch system in cultured neurons from B6.Cg-Tg (Fos-tTA,Fos-shEGFP) mice (**Fig. S7, A** to **C**), each well of a 24-well glass-bottom plate received 100 ng of the structural monomer plasmid (*UbC*-CytoTape-HA), 20 ng of the timestamp monomer plasmid (*UbC*-CytoTape-HaloTag), 25 ng of the GFP nanobody-bearing monomer plasmid (*UbC*-CytoTape-OLLAS-GFPnanobody), with (**Fig. S7C**) or without (**Fig. S7B**) 50 ng of the backfiller plasmid (*UbC*-Myc-EGFP(dark)), and pUC19 plasmid. pUC19 plasmid was supplemented to bring the total amount of plasmid DNA to 1,500 ng per transfection.

For experiments related to the GFP nanobody-based CytoCatch system in cultured neurons from wild-type Swiss Webster mice (**Fig. S7D**), each well of a 24-well glass-bottom plate received 100 ng of the structural monomer plasmid (*UbC*-CytoTape-HA), 20 ng of the timestamp monomer plasmid (*UbC*-CytoTape-HaloTag), 25 ng of the GFP nanobody-bearing monomer plasmid, 50 ng of the backfiller plasmid (*UbC*-Myc-EGFP(dark)), 25 ng of the signal monomer plasmid (*Fos*-mEGFP), and 1280 ng of pUC19 plasmid. For experiments related to CytoCatch-Spy for abundance-resolved molecule archiving (**Fig. 5**), each well of a 24-well glass-bottom plate received 100 ng of the structural monomer plasmid (*UbC*-CytoTape-HA), 20 ng of the timestamp monomer plasmid (*UbC*-CytoTape-HaloTag), 25 ng of the CytoCatch-Spy monomer plasmid (*UbC*-CytoTape-FLAG-SpyCatcher003), 50 ng of the backfiller plasmid (*UbC*-SpyTag003-sfGFP(dark)-FLAG), 25 ng of the signal monomer plasmid (*Fos*-SpyTag003-sfGFP(dark)-V5), and 1280 ng of pUC19 plasmid. After a 45-min incubation with DNA-calcium phosphate precipitates, cells were washed with a preformulated acidic buffer, HMEM (made by MEM buffer supplemented with 15 mM HEPES and then adjusted to a final pH of 6.70−6.74 with acetic acid (Millipore Sigma) over the time course of 6 h) to remove residual precipitates. Cells were then placed back in the cell culture incubator.

### CyTRACE expression in U2OS cells

U2OS cells at a low passage number (<10 passages) were plated at 40% confluence onto a 200-mm tissue culture dish.

Dulbecco’s Modified Eagle’s Medium (DMEM; Gibco; containing high glucose, GlutaMAX supplement, and pyruvate) supplemented with 10% HI FBS (Corning) and penicillin/streptomycin (Gibco) was used as the U2OS cell culture medium. Cells were then grown in a cell culture incubator at 37 ^◦^C with 5% CO_2_. After reaching 50-70% confluence, cells were transferred to 24-well glass-bottom plates via trypsin treatment. The 24-well glass-bottom plates were pre-treated with 70 µL Matrigel in the cell culture incubator for 40 min before cell plating. 24 h after cell plating, genes were delivered using the *Trans*IT-X2 Dynamic Delivery System kit (Mirus Bio). For experiments related to screening structural monomer designs in U2OS (**Fig. S1**), each well of a 24-well glass-bottom plate received 100 ng of one of the candidate structural monomer plasmids (*UbC*-CytoTape-HA; *UbC*-CytoTape-vivo-HA; *UbC*-XRI-HA; *UbC*-1POK (E239Y)-GG-MBP-HA), 20 ng timestamp monomer plasmid (*UbC*-CytoTape-HaloTag), and 180 ng pUC19 plasmid. For experiments related to CytoSense-NS3pro in U2OS (**Fig. 2, F** to **M**), each well of a 24-well glass-bottom plate received 100 ng structural monomer plasmid (*UbC*-CytoTape-vivo-HA), 20 ng timestamp monomer plasmid (*UbC*-CytoTape-HaloTag), 10 ng CytoSense-NS3pro monomer plasmid (*UbC*-CytoTape-V5-SMASh), and 170 ng pUC19 plasmid. For experiments related to CytoCatch-Spy in U2OS (**Fig. 6**), each well of a 24-well glass-bottom plate received 100 ng structural monomer plasmid (*UbC*-CytoTape-vivo-HA), 20 ng timestamp monomer plasmid (*UbC*-CytoTape-HaloTag), 20 ng CytoCatch-Spy monomer plasmid (*UbC*-CytoTape-FLAG-SpyCatcher003), 20 ng SpyTag-FT monomer plasmid (*UbC*-SlowFT-FLAG-SpyTag003), and 140 ng pUC19 plasmid. Plasmids were first diluted in 50 µL Opti-MEM medium, followed by the addition of 0.9 µL *Trans*IT-X2 reagent. The mixture was incubated at room temperature for 25 min, and then added to every single well.

### CyTRACE timestamp labeling

CyTRACE was labeled with cell-permeant HaloTag-ligand Janelia Fluor (JF) dyes (Promega) (**Fig. 2, F** to **M** and **Fig. 3**, **N** to **O** and **Fig. 4, H** to **N** and **Fig. 5** and **Fig. 6**). JF_503_, JF_585_, and JF_635_ were used in this study for timestamps. The JF dyes were in lyophilized powder form and stored at -20 °C before use. The JF dye powder was first dissolved in 50 µL dimethyl sulfoxide (DMSO) and then diluted into 10 mL of U2OS cell culture media (for U2OS cells) or into 10 mL of neuronal culture media (for cultured neurons) at a final concentration of 0.1 µM. The dyed medium was filtered using a sterile 0.22 µm syringe filter before being added to the culture. It was then used to replace the original media in cell cultures at designated time points. During the dye-switching process, the medium containing the original dye was fully removed, followed by thorough washes of the cell cultures three times with 37 °C DMEM for U2OS cells, and MEM for cultured neurons. Finally, the cells in each well of the 24-well plate were cultured in 500 µL of fresh U2OS cell culture media (for U2OS cells) or 1 mL neuronal culture media (for cultured neurons) supplemented with a new JF dye at 0.1 µM final concentration in the cell culture incubator at 37 °C with 5% CO_2_.

### CyTRACE recording and chemical stimulation in cultured neurons

For CytoSense-NS3pro related experiments (**Fig. 2, B** to **E**), 10 mM Asunaprevir (ASV) dissolved in DMSO (MedChemExpress BMS-650032) was stored at -80 °C before use. 4 days after neuron calcium phosphate transfection, 10 mM ASV stock solution was diluted in 500 µL neuronal culture medium to a final concentration of 3 µM, and the original medium in each well was replaced with the ASV-containing medium in each well in the 24-well plates. ASV treatment lasted for 3 days, until day 7 post-transfection, when the cells were fixed and immunostained for imaging. For CytoSense-mTORC1/S6K experiments, neurons were stimulated according to the post-transfection time points, drug concentrations, and treatment durations described for each experiment in the Results section. Rapamycin (MedChemExpress HY-10219) was stored as a 10 mM stock solution dissolved in DMSO at –80 °C before use.

Everolimus (MedChemExpress HY-10218) was stored as a 10 mM stock solution dissolved in DMSO at -80 °C before use. Torkinib (MedChemExpress HY-10414) was stored as a 10 mM stock solution dissolved in DMSO at –80 °C before use. For specific experiments, rapamycin was used at either 25 nM or 50 nM (**Fig. 3**), torkinib at 1 μM (**Fig. 3, N** to **P**, concentration indicated in Results and figures), and everolimus at 50 nM (**Fig. 3, N** to **P**). Drugs were diluted from stock solutions into neuronal culture medium to a final volume of 500 μL. Before drug treatment, the original culture medium from neuron cultures was transferred into a fresh 24-well plastic-bottom plate, where the medium from different neuron cultures was stored in separate wells and kept in the neuronal incubator until the end of the drug stimulation. Cells were incubated with the drug-containing medium for the durations indicated in the Results section. After treatment, the drug-containing medium was removed, and cells were washed three times with MEM at 5 min intervals. The original conditioned neuronal culture medium was then returned to the cells for subsequent culture at 37 °C with 5% CO₂. For forskolin (FSK) stimulation (**Fig. 5** and **Fig. S7**), FSK powder was dissolved in DMSO to prepare a 5 mM stock solution, which was stored at -20 °C. Then, an FSK stimulation medium was prepared by mixing the 5 mM FSK solution with fresh neuronal culture medium for a final concentration of 30 µM. The FSK stimulation medium was filtered using a sterile 0.22 µm syringe filter (VWR, Avantor) before being added to neurons. The original culture medium from neuron cultures was transferred into a fresh 24-well plastic-bottom plate, where the medium from different neuron cultures was stored in separate wells and kept in the neuronal incubator until the end of the FSK stimulation. Then, 500 µL of FSK stimulation medium was added to each well containing neuron cultures. The neuron cultures were placed back into the incubator and incubated for 1 h. Finally, the FSK stimulation medium was removed, and wells were washed three times with prewarmed MEM, and the original neuronal culture medium was transferred back into the corresponding wells. The neuron cultures were then returned to the cell culture incubator.

### Chemical stimulation for CytoSense-NS3pro recording in U2OS

For CytoSense-NS3pro experiments in U2OS cells (**Fig. 2, F** to **M**), Asunaprevir (ASV; MedChemExpress, BMS-650032) was dissolved in DMSO to prepare a 10 mM stock solution and stored at −80°C before use. Before treatment, the 10 mM ASV stock was diluted into U2OS culture medium to a final concentration of 3 μM. The original culture medium was removed, and 500 μL of ASV-containing medium was added to each well. Cells were treated for the durations indicated in the Results section. After treatment, the drug-containing medium was removed, and cells were washed three times with fresh DMEM before fresh U2OS culture medium was added and cells were returned to the incubator.

### Antibodies and cell morphology staining

Primary antibodies (1:1000 for immunofluorescence of cultured cells): anti-V5 (Invitrogen R960-25), anti-FLAG (Invitrogen 740001), anti-HA (Cell Signaling Technology 3724; Invitrogen 26183), anti-pS6 (Cell Signaling Technology 2211). Fluorescent secondary antibodies (1:1000 for immunofluorescence of cultured cells): Goat anti-Mouse IgG1 Alexa Fluor 488 (Invitrogen A-21127), Goat anti-Mouse IgG2a Alexa Fluor 488 (Invitrogen A-21131), Goat anti-Mouse IgG2a Alexa Fluor 546 (Invitrogen A-21133), Goat anti-Rabbit IgG(H+L) Alexa Fluor Plus 488 (Invitrogen A-32731), and Goat anti-Mouse IgG1 Alexa Fluor 647 (Invitrogen A-21240).

Nissl stain: NeuroTrace 435/455 Blue Fluorescent Nissl Stain (Invitrogen N21479), 1:1000 for immunofluorescence of cultured cells. Plasma membrane stain: CellMask Deep Red Actin Tracking Stain (Thermo Fisher Scientific, A57245), 1:1000 for immunofluorescence of cultured cells. Additional details of primary antibodies, secondary antibodies, as well as HaloTag ligand dyes used in this study are listed in **Supplementary Table S3**.

### Immunostaining of epitope tags in cultured cells

Cells (neurons, U2OS cells) were fixed in 10% buffered formalin (Fisher Scientific) at room temperature (RT) for 10 min, followed by three 5-minute washes in 1×PBS at RT. Blocking was performed in MAXBlock blocking medium (Active Motif) supplemented with 0.1% Triton X-100 and 100 mM glycine for 20 min at RT, followed by three additional 5-min washes in MAXwash washing medium (Active Motif) at RT. Cells were then incubated with primary antibodies diluted in MAXbind staining medium (Active Motif) for 1 h at RT. Afterward, cells underwent three 5-min washes in MAXwash washing medium at RT. Secondary antibodies were applied in MAXbind staining medium and incubated for 1 h at RT. Cells were then washed three times with MAXwash washing medium for 5 min each at RT. Finally, cells were incubated with NeuroTrace 435/455 Blue Fluorescent Nissl Stain (Invitrogen) for 10 min and stored in 1×PBS at 4 °C until imaging.

### Fluorescence microscopy of immunostained cells

Fluorescence microscopy was performed using a spinning disk confocal microscope (Yokogawa CSU-W1 Confocal Scanner Unit on a Nikon Eclipse Ti2 inverted microscope) equipped with a 40× 1.15 numerical aperture water immersion objective (Nikon MRD77410), a 10× objective, a Hamamatsu ORCA-Fusion BT sCMOS camera controlled by the NIS-Elements AR software, and laser/filter sets for 405 nm, 488 nm, 561 nm, and 640 nm optical channels. Multi-channel volumetric imaging was conducted at 0.4 µm per Z-step for each field of view under the 40× objective. Imaging parameters remained consistent across all samples within each experimental set.

### Software for image analysis

Image analysis was performed in ImageJ (National Institutes of Health) and Python.

### Readout information from intensity profiles

Fluorescence intensity profiles along the tape were extracted from images as previously described^14^.

### Time recovery with HaloTag ligand dye-based timestamp

Two HaloTag ligand dyes were used per experiment in this study to recover time axes in neurons and U2OS cells (**Fig. 2, F** to **M**; **Fig. 3, N** to **P**; **Fig. 4, H** to **N**; **Fig. 5, A** to **H**; **Fig. 6, A** to **N**). The methods for identifying JF dye switching points (“timestamps”) and recovering the time axis based on their positions along the tape were performed as previously described^15^.

**Identification of decay onset time and recovery onset time for CytoSense-NS3pro recorded temporal signals.** For the CytoSense-NS3pro recordings (“traces”) under ASV treatment (**Fig. 2, H** to **M**), the decay onset time was defined as the earliest time after the onset time of ASV treatment at which the signal entered a decreasing phase. Traces lacking an identifiable decay onset were excluded from the downstream latency analysis. The interval between the decay onset time and the onset time of ASV treatment was defined as the decay latency. Similarly, the recovery onset time was defined as the time at which the signal reached its minimum following the decay phase. The interval between the recovery onset time and the ASV withdrawal time was defined as the recovery latency.

### Identification of rise onset time for CytoCatch-Spy archived temporal abundance

For the CytoCatch-Spy archived temporal protein abundance dynamics under FSK treatment (**Fig. 5, F** and **H**), the rise onset time was defined as the earliest time after the onset time of FSK treatment at which the abundance entered an increasing phase.

### Time recovery analysis based on CytoCatch-Spy archived fluorescent timers (FT)

For the analysis of the FT-based internal timer, recordings from the 46 h and 64 h experimental groups were analyzed separately (**Fig. 6, D** to **I**). For each trace, normalized red and blue FT intensity profiles were obtained by dividing each channel by its maximum intensity along the trace (**Fig. 6, D** and **F**). The red-to-blue ratio profile was then calculated along the tape (**Fig. 6, D** and **F**). Next, the cumulative fraction profile was calculated as the cumulative integral of the red-to-blue ratio profile normalized by its total integral across the time range recovered by HaloTag timestamp (**Fig. 6H**). The mean cumulative fraction profile across all traces within the same experimental group was defined as the calibration curve. This calibration curve was then used to recover time from FT profiles. The absolute error between FT-recovered time and HaloTag timestamp-recovered time was calculated to assess temporal accuracy for internal timer (**Fig. 6I**).

For the analysis of the FT-based internal clock, recordings from the 46 h and 64 h experimental groups were also analyzed separately (**Fig. 6, J** to **N**). For each trace, the red-to-blue ratio profile was plotted as a function of tape position (**Fig. 6J**). Most traces exhibited a two-segment profile consisting of a central plateau followed by a declining phase toward the tape end. The plateau-to-decline transition point (“transition point”) was defined as the earliest position at which the profile departed from the plateau region and entered the declining phase and was identified for further analysis. The corresponding time of this transition point, recovered using HaloTag timestamping, was defined as the transition time (**Fig. 6L**). The median transition time across all traces within the same experimental group was defined as the calibrated transition time. To construct the internal clock calibration curve, the region between the transition point and the tape end was extracted from each red-to-blue ratio profile (**Fig. S8, B** and **C**). The cumulative fraction profile was calculated within this region and aligned such that all transition points corresponded to the calibrated transition time (**Fig. S8D**). The mean aligned cumulative fraction profile across all traces was defined as the internal clock calibration curve. This calibration curve was then used to recover time between the tape end and transition point for each trace (**Fig. 6M**). The absolute error between FT clock-recovered time and HaloTag timestamp-recovered time was calculated to assess temporal accuracy of the FT internal clock (**Fig. 6N**).

### Cellular fluorescence intensity analysis for anti-pS6 staining

Multi-channel Z-stacks for the raw confocal images from neurons were imported using the Bio-Formats plugin in ImageJ/FIJI. To account for cellular volume across the Z-axis, a Maximum Intensity Projection was generated for each field of view (FOV), and the channels were split to isolate the Nissl and pS6 signals. Cellular segmentation was performed in the Nissl channel to define Regions of Interest (ROIs). The segmentation pipeline consisted of noise reduction with a median filter, Otsu automated thresholding, watershed transformation, and particle analysis by size and circularity. The resulting ROIs were mapped onto the pS6 staining channel. The Mean Gray Value for pS6 was recorded for every identified cell with batch processing across FOVs and experimental conditions, and exported for secondary statistical analysis. Background intensity was measured from the dimmest region of a randomly selected FOV for each experimental time point, calculated separately for control and drug-treated conditions. The averaged background intensity for each respective group was subtracted from the raw pS6 mean gray values during statistical analysis (**Fig. 4, D** and **E**). For the physiological sensitivity analysis, neurons from the no-rapamycin control group at the final sampling time point were classified according to their anti-pS6 fluorescence intensity. To define the low-pS6 intensity range, cellular anti-pS6 intensities from rapamycin-treated neurons at the same time point were used as the reference. A one-dimensional Mahalanobis distance distribution relative to the rapamycin-treated population mean and covariance was calculated. The 95th percentile of the within-rapamycin-treated Mahalanobis distance distribution was used as the initial boundary. Because the rapamycin-treated anti-pS6 distribution had a narrow variance, this boundary was expanded by a factor selected from the first stable plateau in the number of control neurons included as the expansion factor was swept. Using this expanded Mahalanobis-distance cutoff, control neurons falling within the anti-pS6 range were classified as low anti-pS6 cells, whereas the remaining control neurons were classified as high anti-pS6 cells.

### Statistical analysis

Statistical analysis was performed in Prism (GraphPad) and Python. All statistical tests were two-sided.

## Acknowledgments

We thank Jiahui Ding, Diane Fingar, Yongjie Hou, William Joesten, Dongqing Shi, Yuwei Tang, and Wenjing Wang for the discussion. Y.Y. acknowledges the Rackham Graduate Student Research Grant. L.Z. acknowledges the Michigan Neuroscience Institute Postdoctoral Advancement Program and the eLife Ambassadors Program. C.L. is supported by the National Institutes of Health (NIH) Director’s New Innovator Award (DP2MH140133), the Glenn Foundation for Medical Research and American Federation for Aging Research Grant Award for Junior Faculty, and the Klingenstein Fellowship Award in Neuroscience.

## Author contributions

Y.Y. and C.L. conceived the concept of CyTRACE and made high-level plans for this project. Y.Y. designed the constructs and performed cell culture experiments with the help and input from L.Z., Y.W., and C.L. Y.Y., L.Z. prepared U2OS cells with the help from J.L. J.L. prepared cultured neurons. Y.Y., Y.W. analyzed the data with the help from L.Z. and C.L. Y.Y., Y.W., L.Z., and C.L. interpreted the data. Y.Y. and C.L. wrote the manuscript with the help from L.Z., Y.W., N.X. C.L. supervised the project.

## Data availability

Plasmids and the corresponding sequence maps of constructs reported in this paper will be available at Addgene upon completion of the deposition process. Sequences will also be available at GenBank upon completion of the deposition process. The dataset is available at Zenodo (https://doi.org/10.5281/zenodo.20603086).

## Code availability

The code used in this work for data analysis is available at GitHub (https://github.com/LinghuLab/CyTRACE).

## Supplementary files

Supplementary Table S1 includes sequences of protein motifs used in this study. Supplementary Table S2 includes plasmid constructs used in cell culture in this study. Supplementary Table S3 includes reagents and resources used in this study.

---

**Fig. S1.**
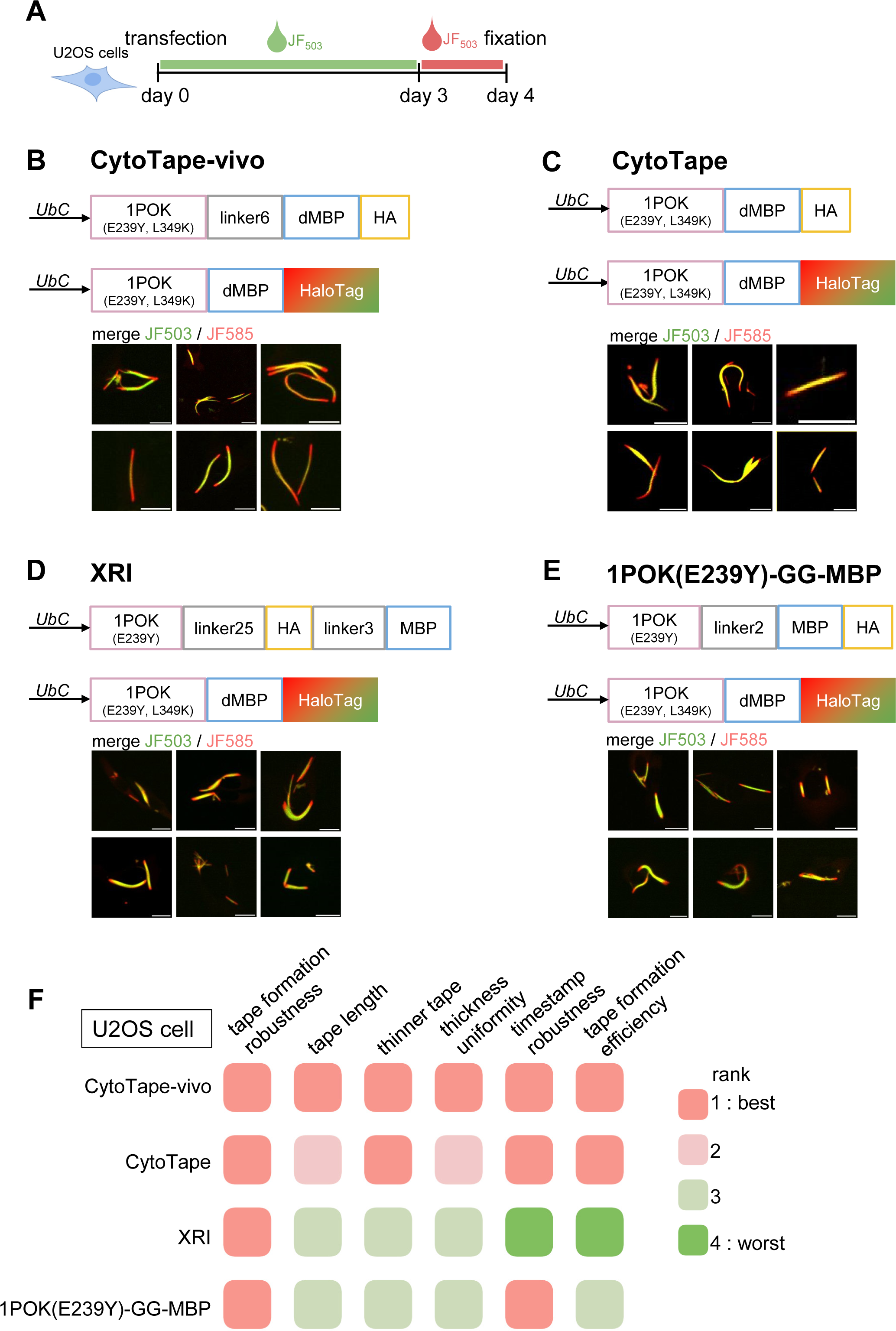
Characterization of protein assemblies from distinct structural monomer designs in U2OS cells. (**A**) Experimental timeline for characterization of distinct structural monomer designs with HaloTag-based timestamps in U2OS cells. (**B-E**) Construct schematics and representative confocal images of CytoTape-vivo, CytoTape, XRI, and 1POK(E239Y)-GG-MBP protein assemblies in U2OS cells. Scale bars, 25 μm. (**F**) Comparison of tape quality metrics across structural monomer designs after 4 days of expression in U2OS cells, including tape formation robustness (fraction of assemblies that have linear, tape-like morphology rather than puncta or short fragments); tape length (length of tapes); thinner tape (smaller tape thickness); thickness uniformity (consistency of tape thickness along the tape), timestamping robustness (fraction of tapes exhibiting clear timestamps from HaloTag ligand dye switching), and tape formation efficiency (the number of tapes per imaging field of view in U2OS cell cultures).

**Fig. S2.**
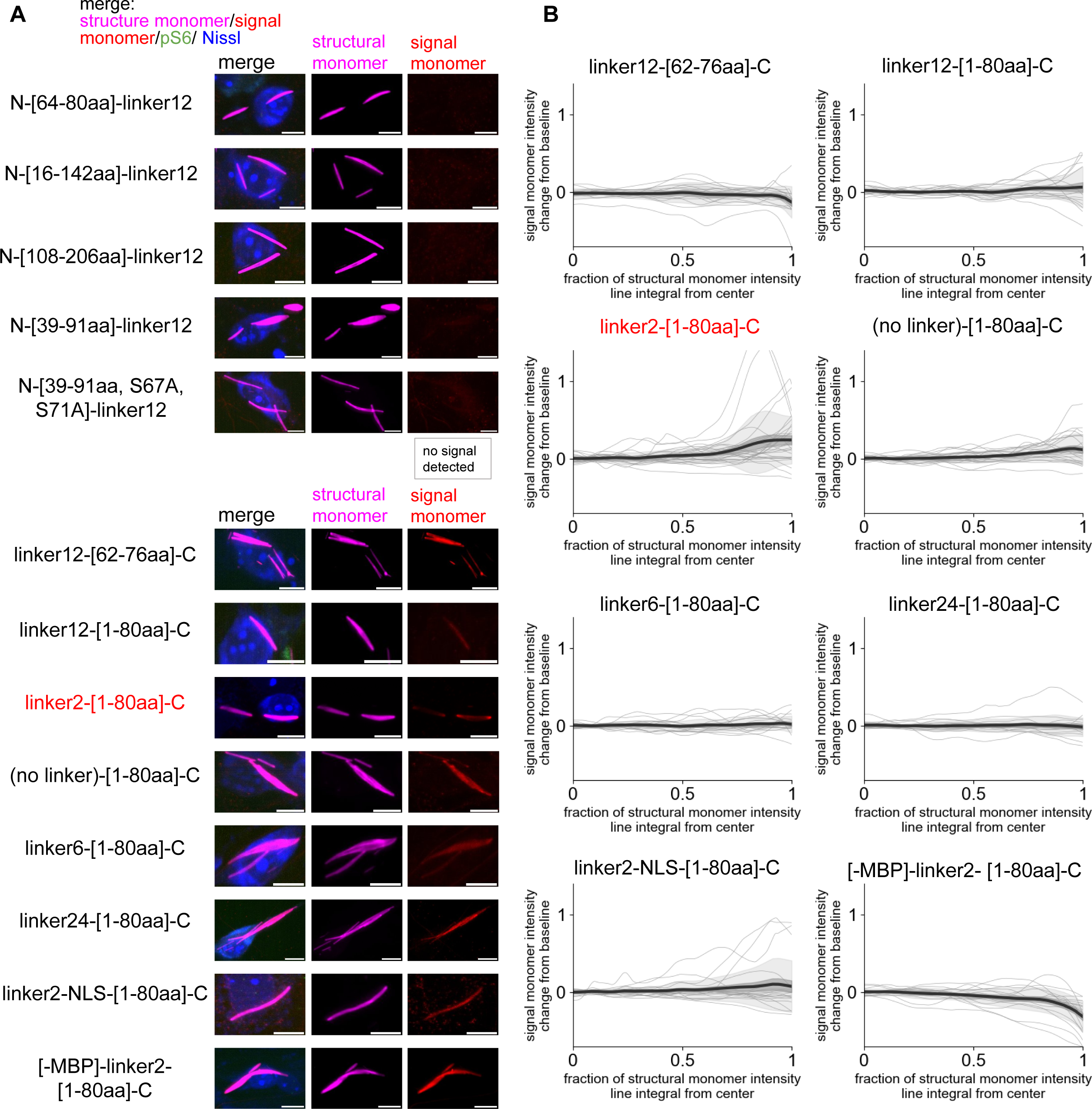
Characterization of candidate CytoSense-mTORC1/S6K signal monomer designs. Cultured neurons were treated with 25 nM rapamycin for 3 h at 7 days after transfection of the same structural monomer (CytoTape) and distinct signal monomer designs, followed by fixation and immunofluorescence imaging. (**A**) Representative confocal images of tapes from each design group. In a subset of design groups, the presence of the corresponding signal monomer variant increased the tape thickness at the center of tape, suggesting that those variants promoted the unwanted lateral monomer binding on the tape. aa, amino acid. Scale bars, 10 µm. (**B**) Population analysis of signal monomer intensity change from baseline profiles plotted as a function of the fraction of the line integral of the structural monomer intensity, for signal monomer variants that had detectable expression. Black center line, mean; darker shaded boundary, s.e.m.; lighter shaded boundary, s.d.; gray thin lines, data from individual tapes. *n* = 15 (linker12-[62–76aa]-C), 18 (linker12-[1–80aa]-C), 23 ((no linker)-[1–80aa]-C), 26 (linker2-[1–80aa]-C), 18 (linker6-[1–80aa]-C), 18 (linker24-[1–80aa]-C), 20 (linker2-NLS-[1–80aa]-C), and 19 ([-MBP]-linker2-[1–80aa]-C) tapes from the same numbers of cells, respectively.

**Fig. S3.**
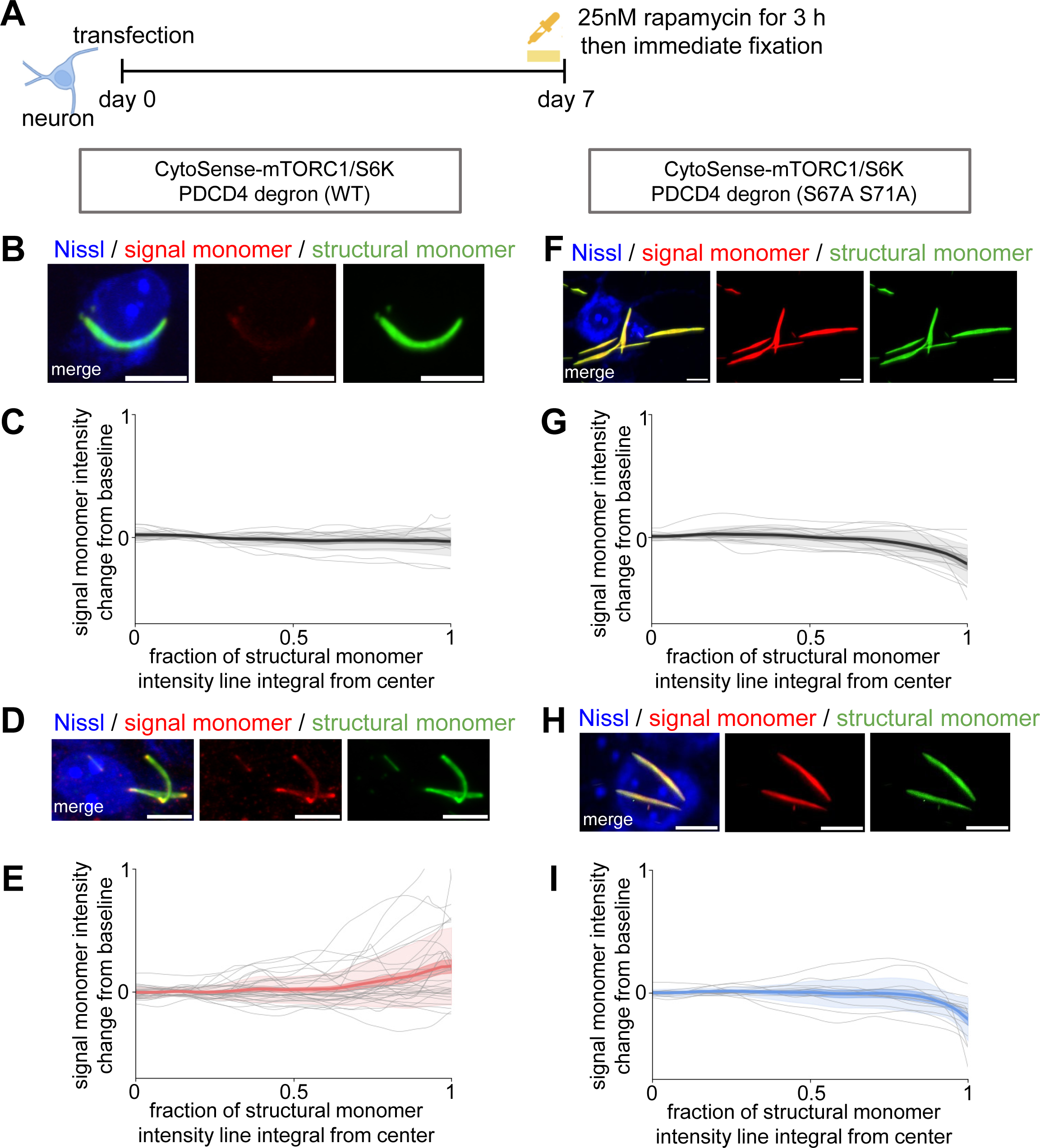
Inactivation of the PDCD4 degron via mutation abolishes CytoSense’s sensitivity to mTORC1–S6K signaling. (**A**) Experimental timeline. Cultured neurons were treated with 25 nM rapamycin for 3 h at 7 days after transfection of the same structural monomer (CytoTape) and CytoSense-mTORC1/S6K signal monomer with or without the S67A and S71A mutations in the PDCD4 degron, followed by immediate cell fixation. (**B-E**) CytoSense-mTORC1/S6K with unmutated (WT) PDCD4 degron. Representative confocal images of tapes in no-rapamycin control (**B**) and rapamycin-treated (**D**) groups. Scale bar, 10 µm. Population analysis of signal monomer intensity change from baseline plotted as a function of the fraction of the structural monomer intensity line integral, in no-rapamycin control (**C**, gray, *n* = 13 tapes from 13 cells) and rapamycin-treated (**E**, red, *n* = 31 tapes from 31 cells) groups. (**F-I**) CytoSense-mTORC1/S6K with mutated PDCD4 degron (S67A, S71A). Representative confocal images of tapes in no-rapamycin control (**F**) and rapamycin-treated (**H**) groups. Scale bar, 10 µm. Population analysis of signal monomer intensity change from baseline plotted as a function of the fraction of the structural monomer intensity line integral, in no-rapamycin control (**G**, gray, *n* = 11 tapes from 11 cells) and rapamycin-treated (**I**, blue, *n* = 12 tapes from 12 cells) groups. For (**C**), (**E**), (**G**), (**I**): thick center line, mean; darker boundary, s.e.m.; lighter boundary, s.d.; thin gray lines, data from individual tapes.

**Fig. S4.**
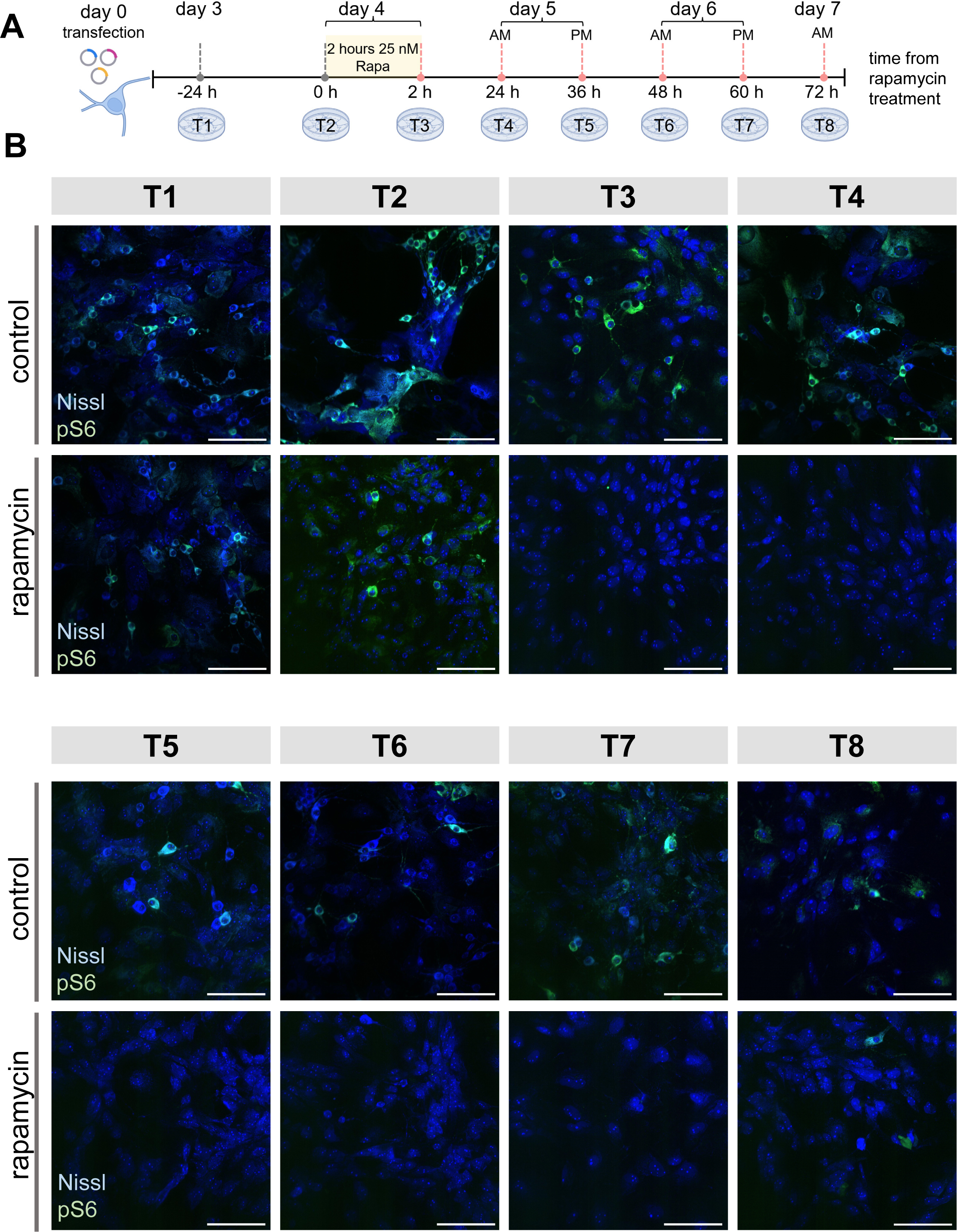
Anti-pS6 (Ser235/236) immunostaining as a ground-truth readout of mTORC1–S6K signaling pathway activity. (**A**) Experimental timeline for time-series sampling of mTORC1–S6K pathway dynamics in cultured neurons, as described in **Fig. 4A**. Cultured neurons expressing the CytoSense-mTORC1/S6K recording system were transiently treated with rapamycin (25 nM, 2 h) at day 4 post-transfection and fixed at distinct time points from day 3 to day 7 (−24 h to 72 h relative to the onset of rapamycin treatment; labeled as T1-T8). (**B**) Representative confocal images of cultured neurons with Nissl staining (blue) and anti-pS6 staining (green) fixed at distinct time points across T1-T8 in no-rapamycin control (top rows) and rapamycin-treated (bottom rows) groups. Scale bars, 100 µm.

**Fig. S5.**
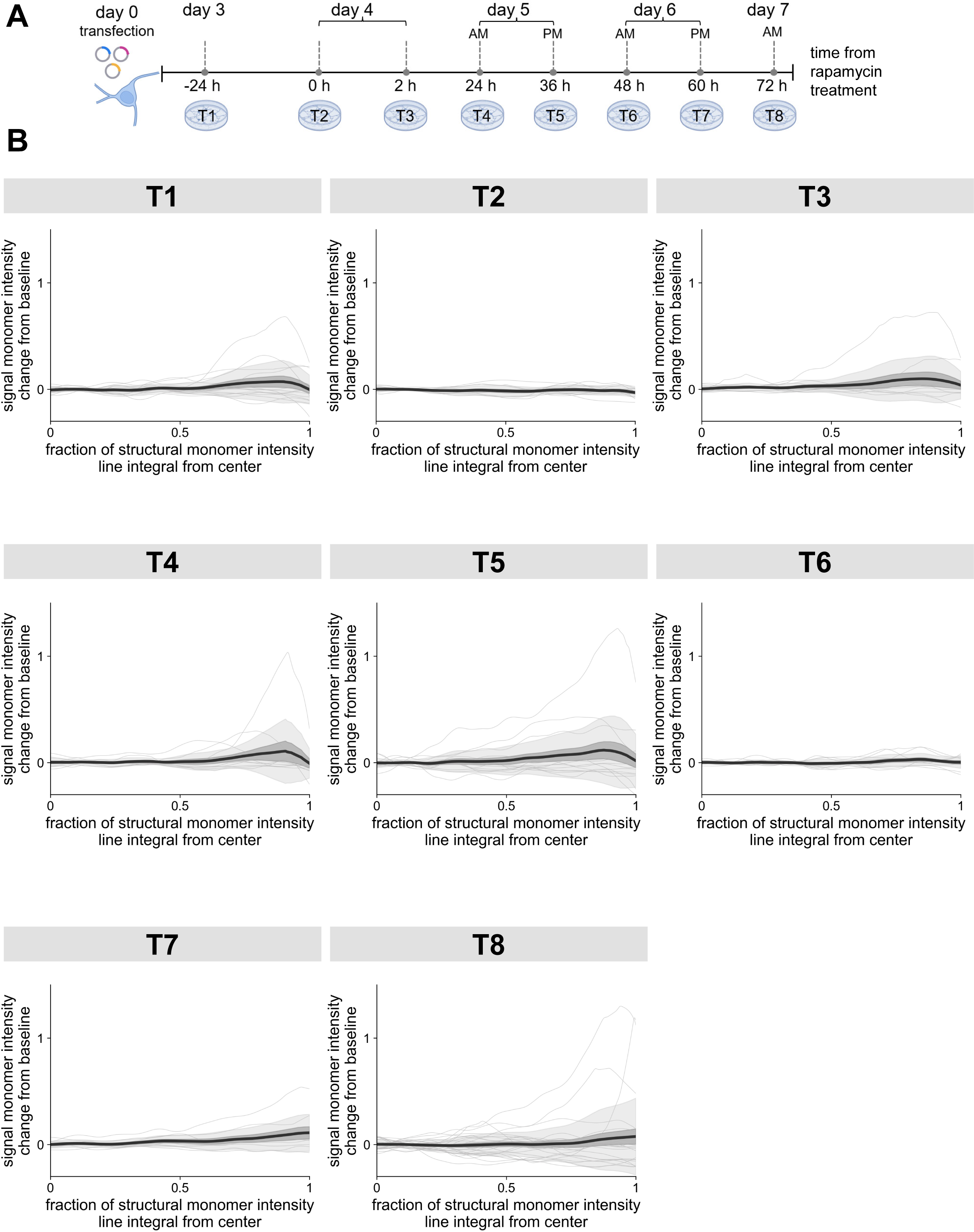
CytoSense-mTORC1/S6K recordings in cultured neurons without pharmacological perturbation. (**A**) Experimental timeline for CytoSense-mTORC1/S6K recording and time-series sampling of mTORC1–S6K pathway dynamics in cultured neurons, as described in **Fig. 4A**. Neurons expressing the CytoSense-mTORC1/S6K recording system were fixed at multiple time points from day 3 to day 7 (−24 h to 72 h relative to the onset of rapamycin treatment for the paired rapamycin-treated group shown in **Fig. 4** and **Fig. S4**; labeled as T1-T8) without rapamycin treatment. (**B**) Population analysis of signal monomer intensity change from baseline plotted as a function of the fraction of the structural monomer intensity line integral, in neurons fixed at distinct time points across T1-T8 in the no-rapamycin control group, under CytoSense-mTORC1/S6K recording. Thick center line, mean; darker shaded boundary, s.e.m.; lighter shaded boundary, s.d.; thin gray lines, data from individual tapes. *n* = 14, 11, 11, 10, 15, 11, 9, 25 tapes for T1-T8, respectively.

**Fig. S6.**
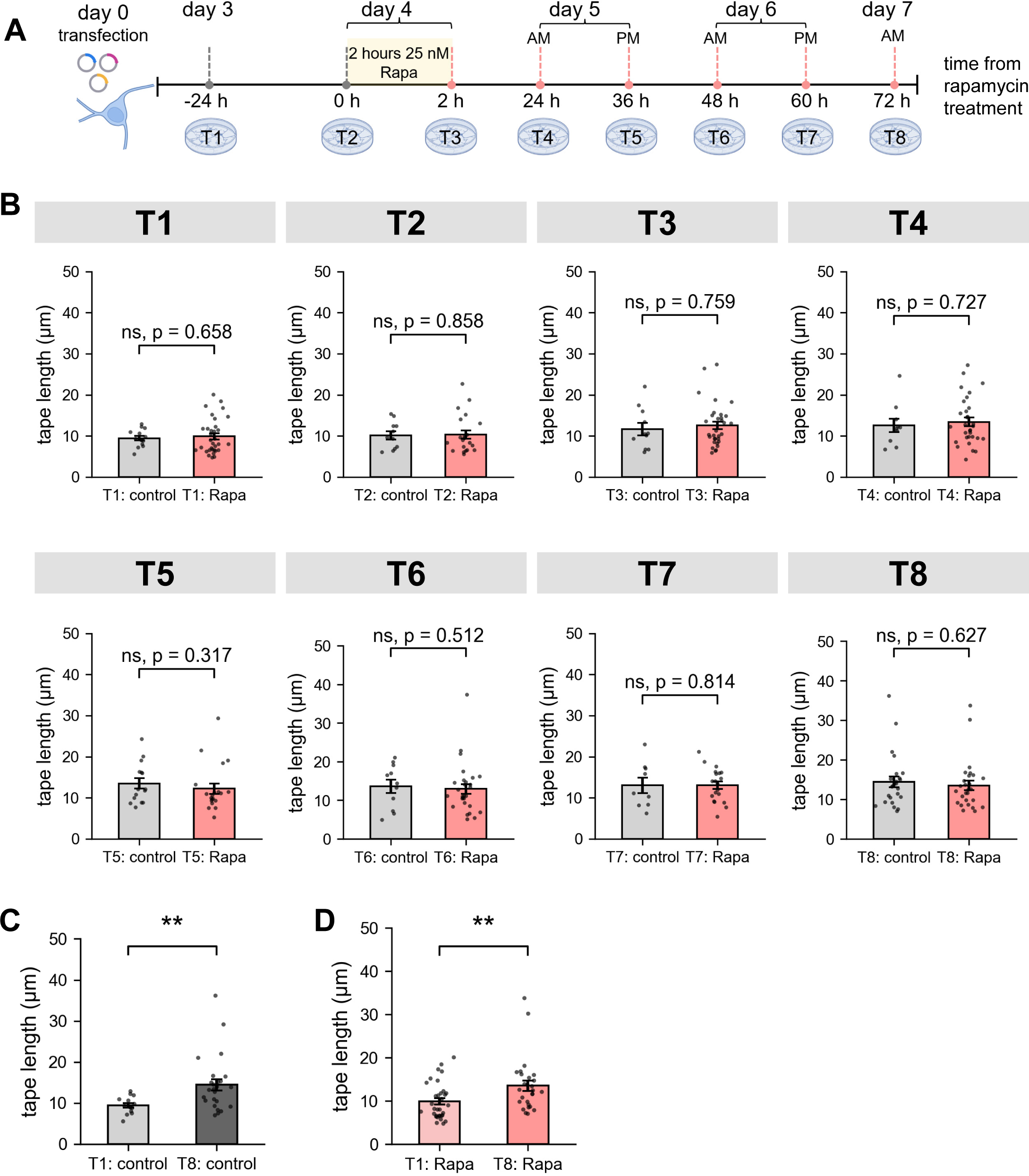
Rapamycin treatment does not significantly affect tape elongation across time. (**A**) Experimental timeline for CytoSense-mTORC1/S6K recording and time-series sampling of mTORC1–S6K pathway dynamics in cultured neurons, as described in **Fig. 4A**. Neurons expressing the CytoSense-mTORC1/S6K recording system were transiently treated with or without rapamycin (25 nM, 2 h) 4 days after transfection and fixed at distinct time points from day 3 to day 7 (-24 h to 72 h relative to the onset of rapamycin treatment; labeled as T1-T8). (**B**) Comparison of tape length in no-rapamycin control (gray) and rapamycin-treated (red) groups across T1-T8 time points (mean ± s.e.m.). Each dot represents one tape. Sample numbers of tapes (each tape was from an individual cell) for no-rapamycin control/rapamycin-treated groups at each time point: T1, *n* = 14/33; T2, *n* = 11/21; T3, *n* = 11/32; T4, *n* = 10/31; T5, *n* = 15/20; T6, *n* = 11/28; T7, *n* = 9/20; T8, *n* = 25/27. (**C**) Comparison of tape length between T1 (light gray, *n* = 14 tapes from 14 cells) and T8 (dark gray, *n* = 25 tapes from 25 cells) in the no-rapamycin control group (mean ± s.e.m.). (**D**) Comparison of tape length between T1 (light red, *n* = 33 tapes from 33 cells) and T8 (dark red, *n* = 27 tapes from 27 cells) in rapamycin-treated group (mean ± s.e.m.). Throughout this figure: n.s., not significant (with the exact P values indicated alongside); **, P < 0.01; two-sided Mann-Whitney U test.

**Fig. S7.**
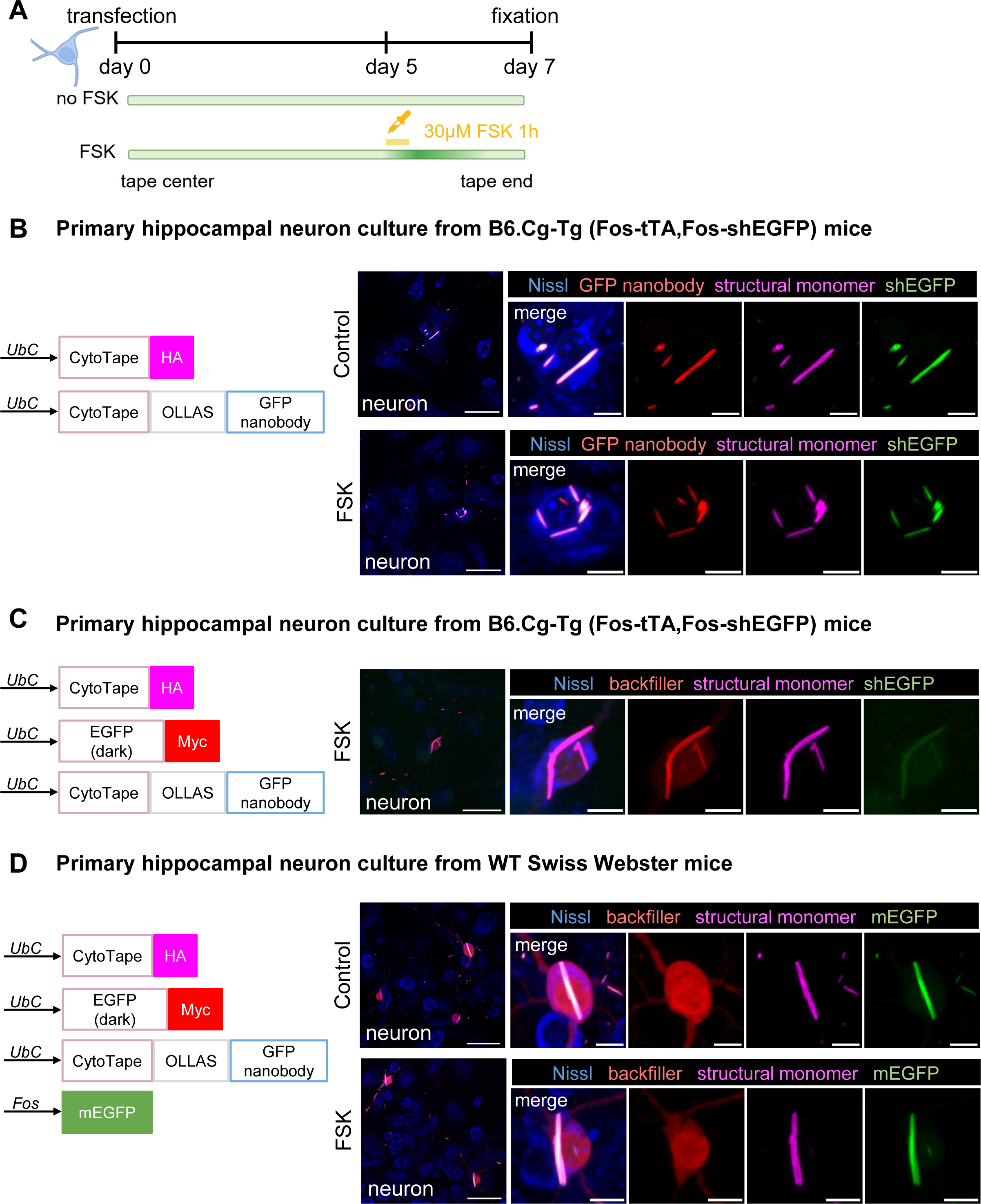
GFP-nanobody-bearing CytoTape monomer archives genomically encoded shEGFP proteins onto the tape but does not resolve their temporal abundance change. (**A**) Experimental timeline (top) and expected density profile of archived shEGFP proteins from tape center to tape end (bottom), using GFP-nanobody-bearing CytoTape monomer to recruit genomically encoded, *Fos* promoter-driven shEGFP as POI in cultured primary hippocampal neurons prepared from the transgenic TetTag mice (B6.Cg-Tg(Fos-tTA,Fos-shEGFP)1Mmay/J). 1 h of 30 μM forskolin (FSK) was applied at day 5 post-transfection and cells were fixed at day 7 post-transfection. (**B** and **C**) Expressing GFP-nanobody-bearing protein tape monomer in neuron cultures from the TetTag transgenic mice. Left, diagrams of transfected constructs, with (**C**) or without (**B**) backfiller co-transfection; right, representative confocal images of formed tapes under no-FSK control or FSK-treated conditions. The shEGFP channel shows the native shEGFP fluorescence, while the structural monomer, GFP nanobody, and backfiller channels show the immunofluorescence for HA, OLLAS, and Myc tags fused to these proteins, respectively. (**D**) Expressing GFP-nanobody-bearing protein tape monomers together with an exogenously introduced *Fos* promoter-driven mEGFP as POI in cultured neurons from wild-type Swiss Webster mice. The green channel shows the native mEGFP fluorescence, while the structural monomer and backfiller channels show the immunofluorescence for HA and Myc tags fused to these proteins, respectively. Scale bars throughout the figure, 50 µm in low-magnification images; 10 µm in zoomed-in tape images.

**Fig. S8.**
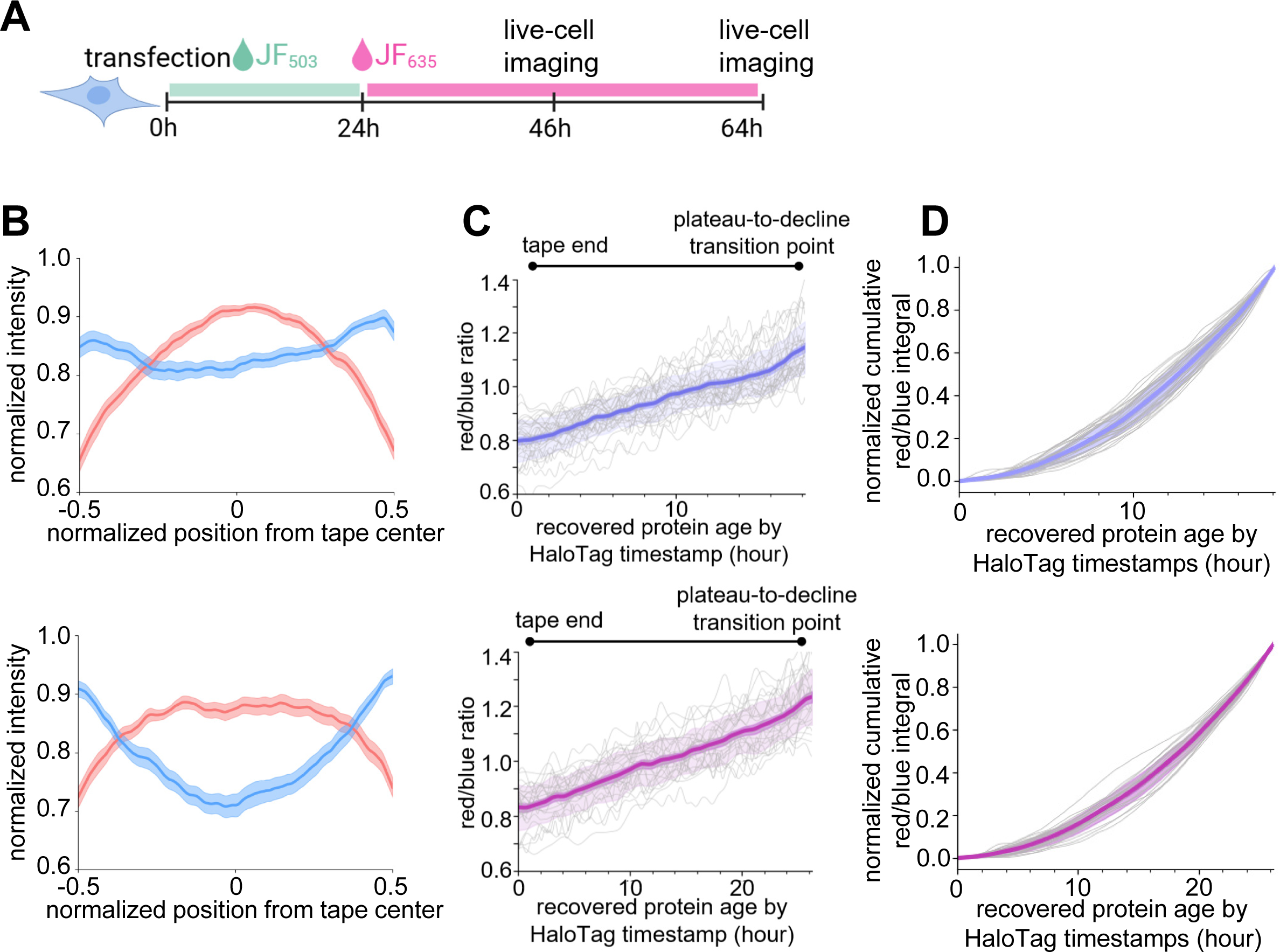
Fluorescent timer proteins archived by CytoCatch-Spy serve as an internal clock for time axis recovery along the tape without timestamps. **(A)** Experimental timeline for CytoCatch-Spy temporal archiving with HaloTag-based timestamps in U2OS cells. Live-cell imaging was performed 46 h (for the 46 h experiment group) or 64 h (for the 64 h experiment group) post-transfection. (**B**) Normalized blue and red fluorescence intensity profiles of archived FTs along the tape for 46 h (top) and 64 h (bottom) experiment groups. Throughout this figure unless specified: center line, mean; dark shaded boundary, s.e.m.; light shaded boundary, s.d.; gray thin lines, data from individual tapes. (**C**) Red-to-blue ratio profiles along the recovered protein age by HaloTag timestamp for the 46 h (top, violet) and 64 h (bottom, purple) experiment groups. Throughout this figure: 46 h experiment group, violet, *n* = 30 tapes from 30 cells; 64 h experiment group, purple, *n* = 32 tapes from 32 cells. (**D**) Normalized cumulative integral of the red-to-blue ratio along recovered time for the 46 h (left, violet) and 64 h (right, purple) experiment groups.

**Supplementary Table S1.**
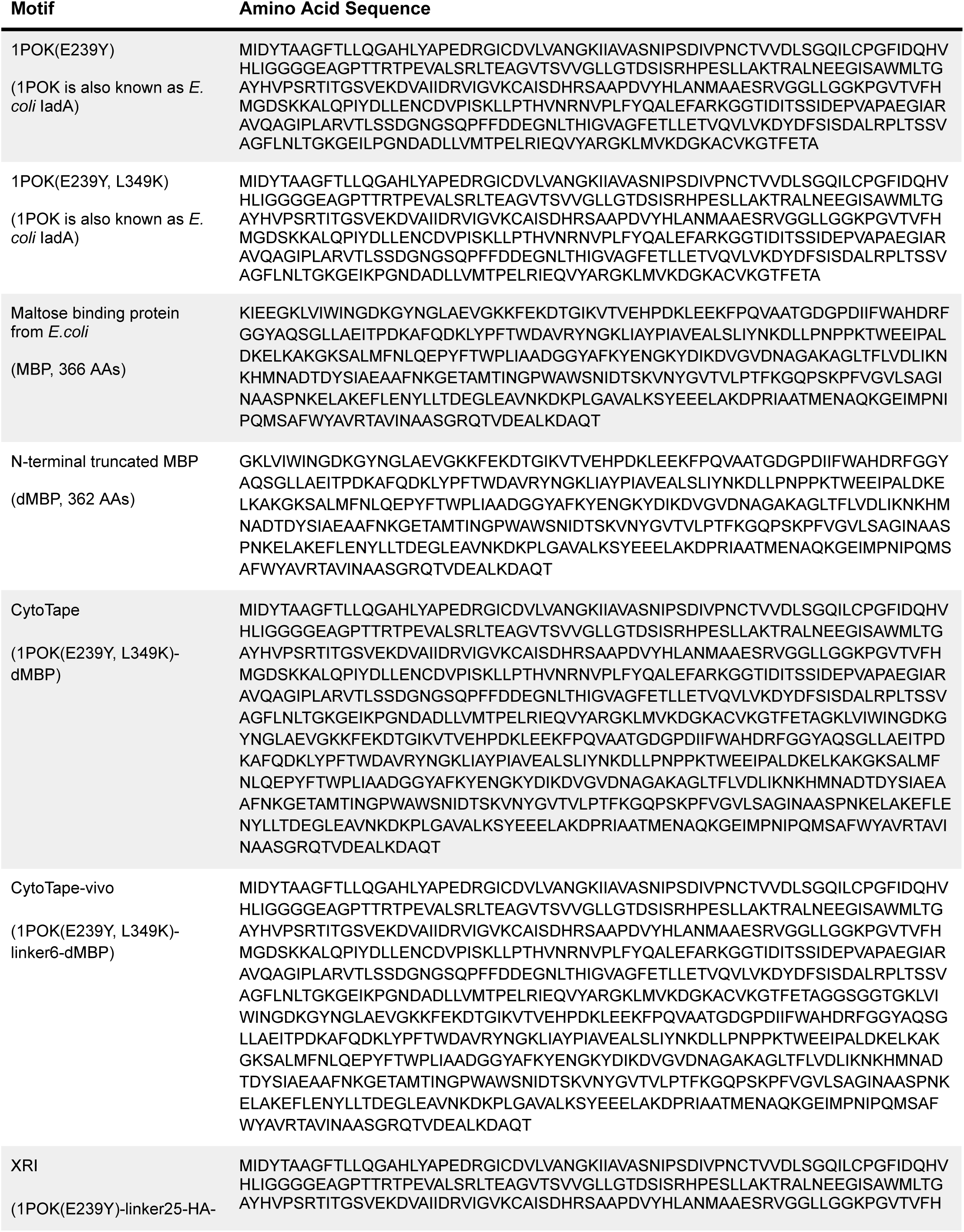
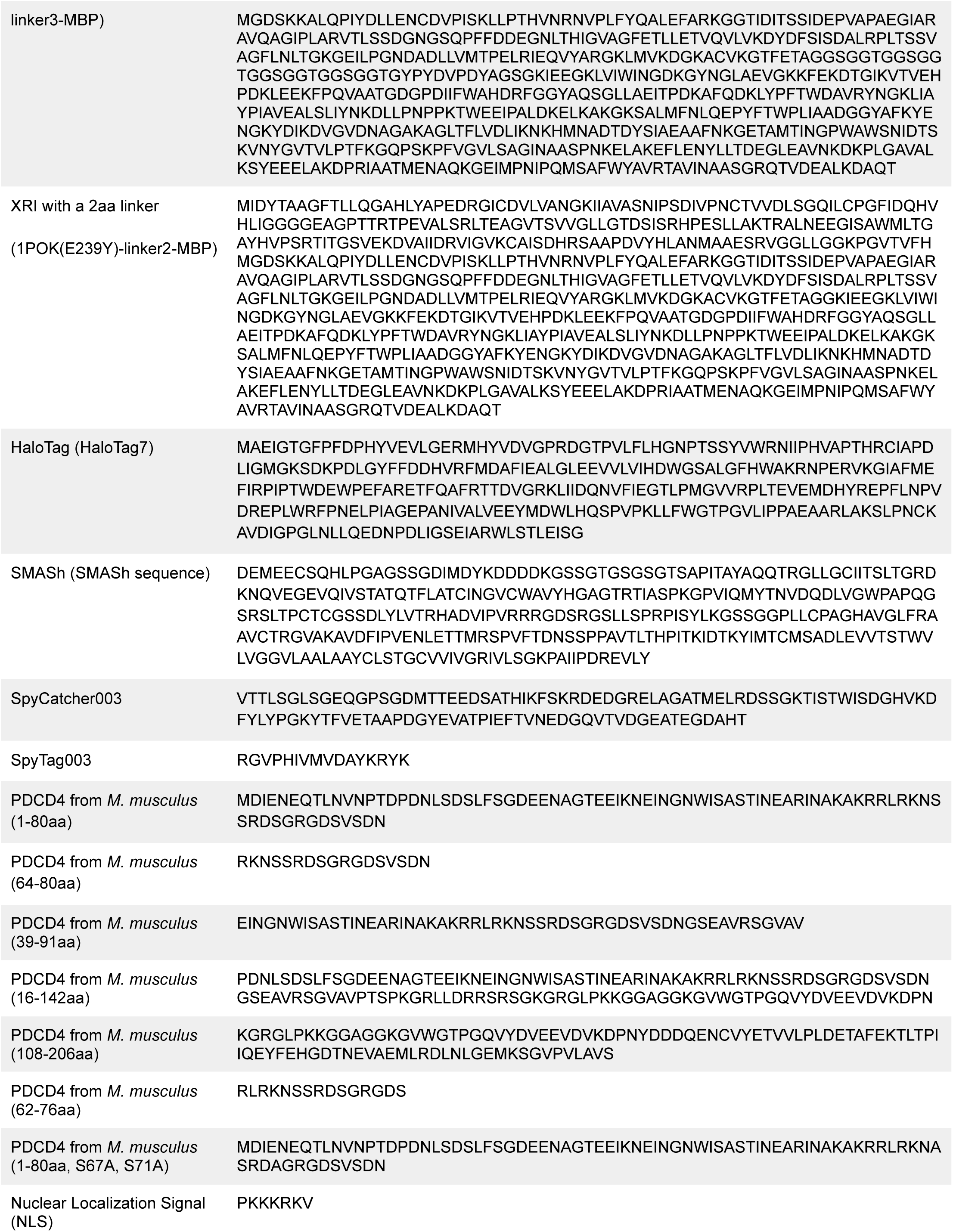
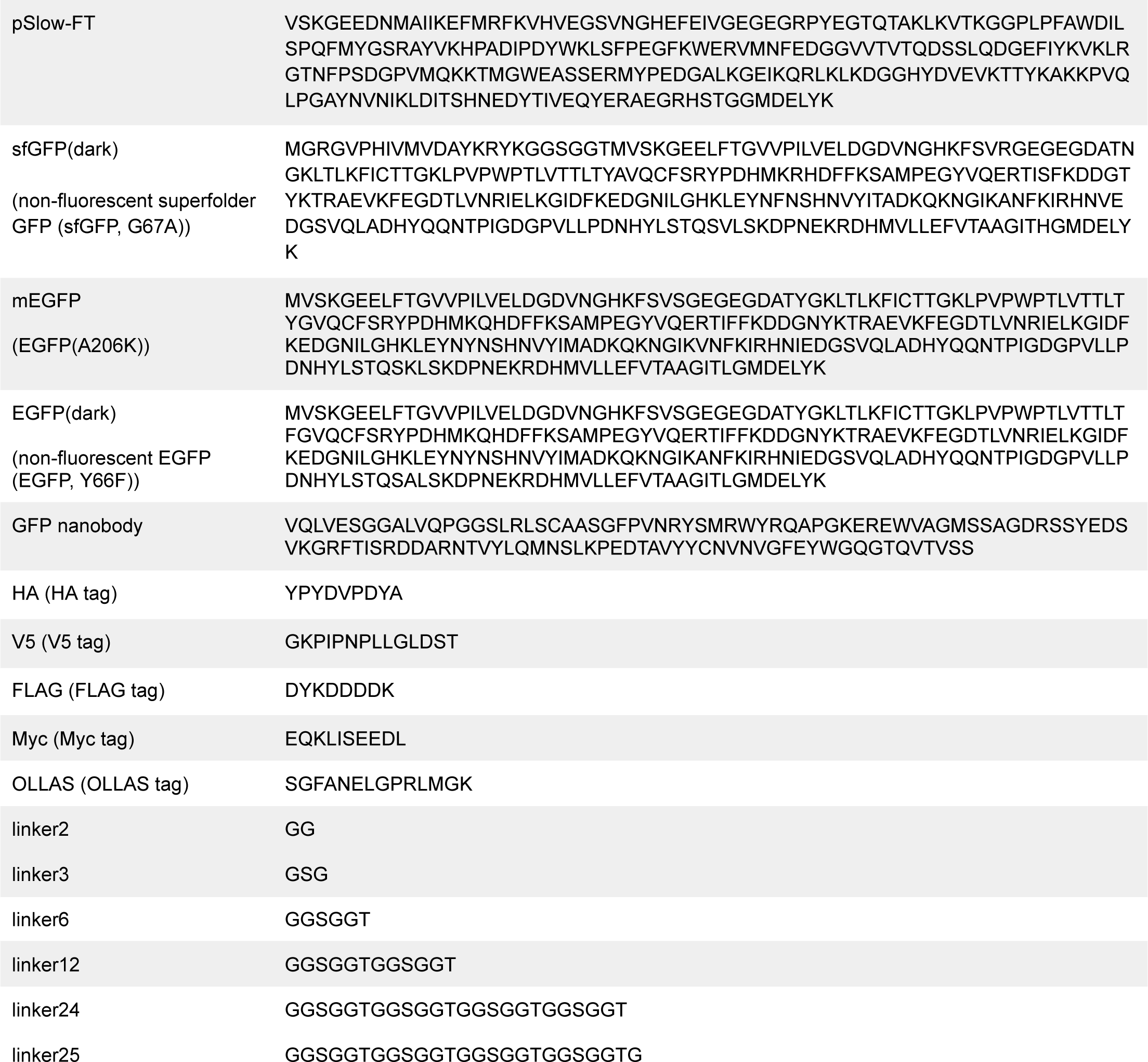
Sequences of protein motifs used in this study.

**Supplementary Table S2.** Constructs used in this study.

| Construct (promoters are underlined) | Tested in |
| --- | --- |
| <u>UbC</u>-CytoTape-HA | Primary neuron cultures |
| <u>UbC</u>-CytoTape-vivo-HA | U2OS |
| <u>UbC</u>-CytoTape-HaloTag | Primary neuron cultures, U2OS |
| <u>UbC</u>-CytoTape-V5-SMASH | Primary neuron cultures, U2OS |
| <u>UbC</u>-PDCD4(64-80aa)-linker12-CytoTape-FLAG | Primary neuron cultures |
| <u>UbC</u>-PDCD4(39-91aa)-linker12-CytoTape-FLAG | Primary neuron cultures |
| <u>UbC</u>-PDCD4(16-142aa)-linker12-CytoTape-FLAG | Primary neuron cultures |
| <u>UbC</u>-PDCD4(108-206aa)-linker12-CytoTape-FLAG | Primary neuron cultures |
| <u>UbC</u>-CytoTape-V5-linker12-PDCD4(62-76aa) | Primary neuron cultures |
| <u>UbC</u>-CytoTape-V5-linker12-PDCD4(1-80aa) | Primary neuron cultures |
| <u>UbC</u>-CytoTape-V5-(no linker)-PDCD4(1-80aa) | Primary neuron cultures |
| <u>UbC</u>-CytoTape-V5-linker2-PDCD4(1-80aa) | Primary neuron cultures |
| <u>UbC</u>-CytoTape-V5-linker6-PDCD4(1-80aa) | Primary neuron cultures |
| <u>UbC</u>-Cytotape-V5-linker24-PDCD4(1-80aa) | Primary neuron cultures |
| <u>UbC</u>-CytoTape-V5-linker2-NLS-PDCD4(1-80aa) | Primary neuron cultures |
| <u>UbC</u>-1POK(E239Y, L349K)-V5-linker2-PDCD4(1-80aa) | Primary neuron cultures |
| <u>UbC</u>-CytoTape-V5-linker2-PDCD4(1-80aa, S67A, S71A) | Primary neuron cultures |
| <u>UbC</u>-Myc-EGFP(dark) | Primary neuron cultures |
| <u>UbC</u>-CytoTape-OLLAS-GFPnanobody | Primary neuron cultures |
| <u>UbC</u>-CytoTape-FLAG-SpyCatcher003 | Primary neuron cultures, U2OS |
| <u>UbC</u>-SpyTag003-EGFP(dark)-V5 | Primary neuron cultures |
| <u>Fos</u>-SpyTag003-mEGFP-Myc | Primary neuron cultures |
| <u>UbC</u>-SpyTag003-sfGFP(dark)-FLAG | Primary neuron cultures |
| <u>Fos</u>-SpyTag003-sfGFP(dark)-V5 | Primary neuron cultures |
| <u>UbC</u>-SlowFT-FLAG-SpyTag003 | U2OS |
| <u>Fos</u>-mEGFP | Primary neuron cultures |

**Supplementary Table S3.**
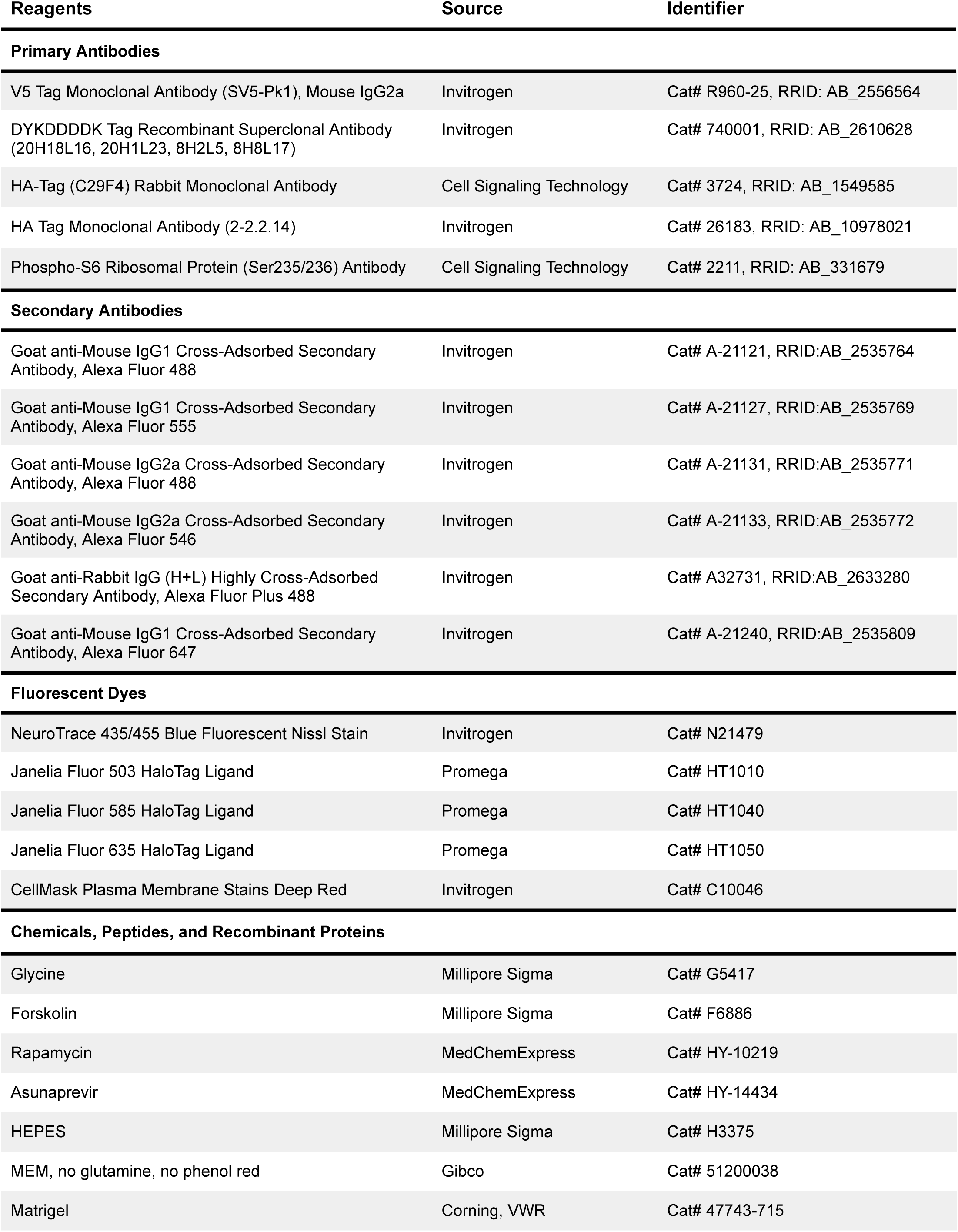
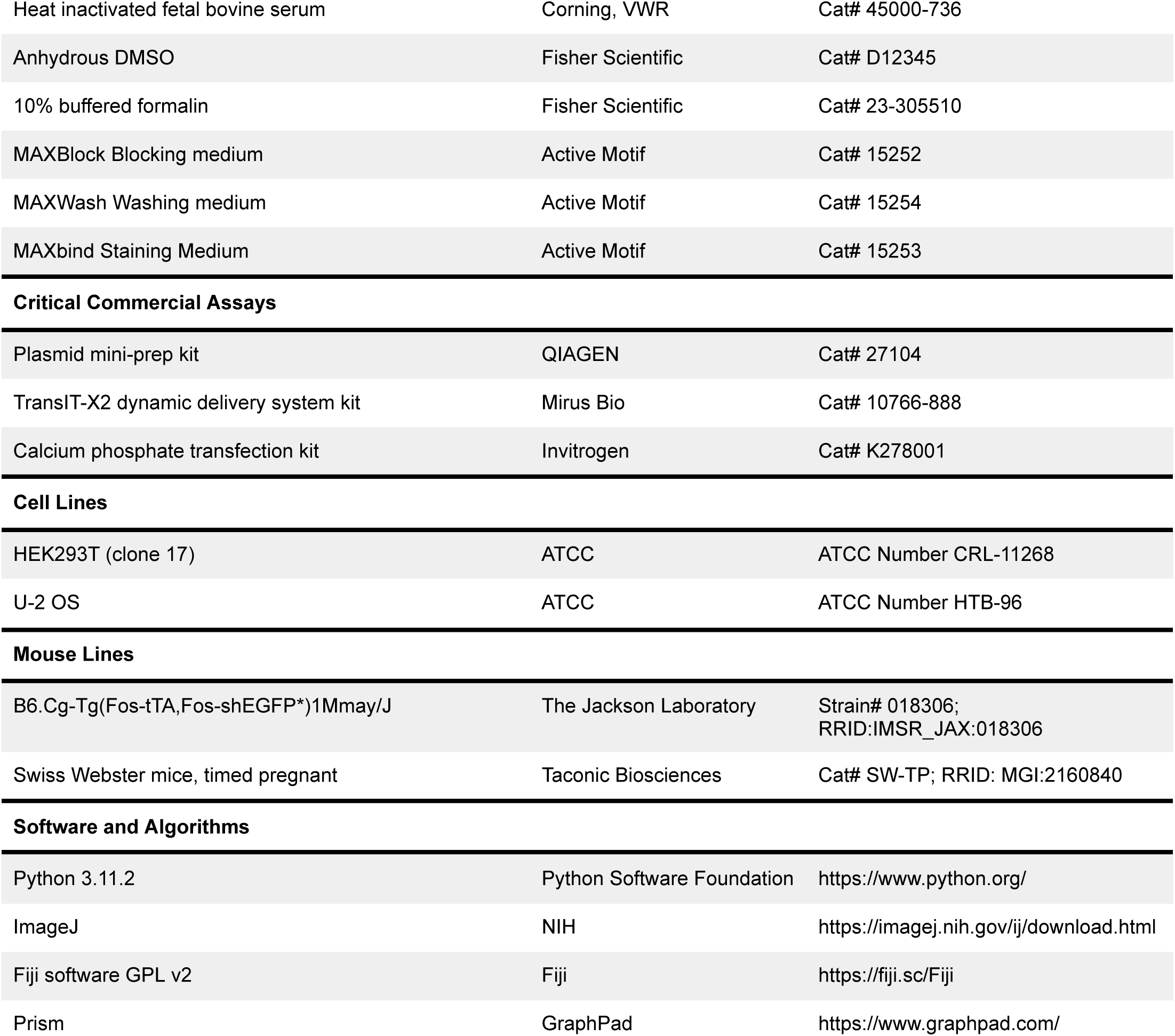
Reagents and resources used in this study

